# Torque-based immune cell chemotaxis in complex environments

**DOI:** 10.1101/2025.11.21.688683

**Authors:** Theresa Jakuszeit, Mathieu Deygas, Mathilde Bernard, Aastha Mathur, Li Wang, Lisa Behrend, Pablo J. Sáez, Pablo Vargas, Raphaël Voituriez, Matthieu Piel

**Affiliations:** Institut Curie and Institut Pierre-Gilles de Gennes, PSL Research University, CNRS, UMR 144, F-75005, Paris, France; Laboratoire Jean Perrin, CNRS, Sorbonne Université, 4 Place Jussieu, 75005 Paris, France; Leukomotion Lab, Paris Cité University, INSERM UMR-S1151, CNRS UMR-S8253, Institut Necker Enfants Malades, F-75015 Paris, France; Cell Communication and Migration Laboratory, Institute of Biochemistry and Molecular Cell Biology, Center for Experimental Medicine, University Medical Center Hamburg-Eppendorf, Hamburg, Germany

## Abstract

Directed migration in chemical gradients is crucial to the immune response, yet how immune cells navigate complex tissues remains incompletely understood. Using in vitro migration assays and theoretical modeling, we uncover distinct chemotactic strategies in two key immune cell types: neutrophils and dendritic cells (DCs). DCs actively steer toward chemokine gradients via a deterministic torque-like reorientation, while neutrophils bias movement by modulating angular noise and speed. A quantitative Fokker–Planck framework decomposes these behaviors into deterministic and stochastic components. Cytoskeletal perturbations show that microtubules enable torque-based navigation in DCs in collagen gels, whereas actomyosin contractility is required for noise modulation employed by neutrophils and DCs in 2D confined migration assays. Despite both achieving directed migration, the two strategies result in opposing macroscopic outcomes: torque-driven cells minimize dispersion, while noise-biased migration enhances population spread. These results reveal distinct navigation aligned with immune function and demonstrate that immune cell chemotaxis is tuned by cytoskeletal architecture and environmental context.

---

In multicellular organisms, cell migration plays a crucial role in a wide range of physiological and pathological processes, from development and tissue repair to immune surveillance and cancer progression [1]. Advances in cell and molecular biology have led to the classification of migrating cells into major subtypes based on their migration mode and associated molecular pathways - such as amoeboid versus mesenchymal, or random versus directed migration [4–6]. Physical models and quantitative *in vitro* studies have been instrumental in identifying key mechanisms of cell migration. They have particularly highlighted the influence of environmental parameters such as adhesion, substrate rigidity, and geometric confinement [6–9]. However, the cellular and physical mechanisms enabling efficient navigation through complex environments - such as tissues - are incompletely understood. In particular, how local cell scale navigation strategies scale up to efficient, macroscopic migration patterns in complex tissues remains an open question.

This question is especially relevant for the immune system, which relies heavily on cell migration. Immune cells represent some of the fastest migrating cells in mammals; traversing complex tissue environments to detect threats or relay information [2, 3]. Despite their diversity, immune cells often share common features: predominantly amoeboid migration, residence in interstitial tissues (e.g., skin) and the ability to respond directionally to molecular cues through chemotaxis. However, their functions can differ markedly. For example, neutrophils and dendritic cells (DCs) are both crucial players in the immune response to infection, injury or abnormal cells, but their direction of migration in response to danger signals is opposite (Fig.1). While neutrophils rapidly converge on sites of infection to neutralize pathogens, DCs undergo a maturation process that includes upregulation of the chemokine receptor CCR7, which guides them to the nearest lymphatic vessel and ultimately to a draining lymph node to initiate an adaptive immune response [10, 11]. Both cell types require efficient chemotactic navigation to fulfil their role in the immune response, yet their tasks - converging on a threat versus migrating away to relay information - are fundamentally different.

**Fig. 1.**
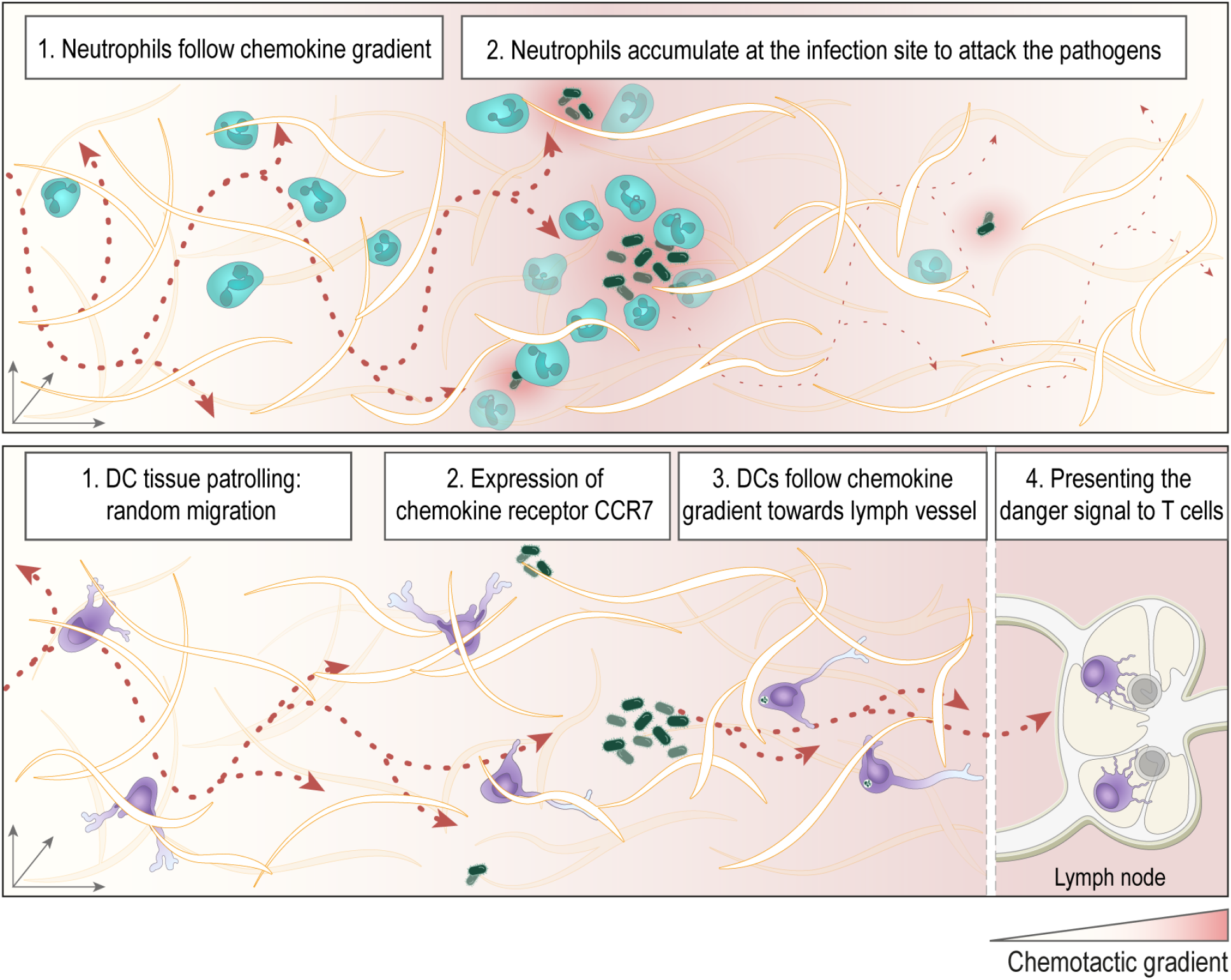
Immune cell chemotaxis in tissues. While neutrophils rapidly converge on sites of infection to neutralize pathogens, DCs undergo a maturation process that includes upregulation of the chemokine receptor CCR7, which guides them to the nearest lymphatic vessel and ultimately to a draining lymph node to initiate an adaptive immune response. Illustration by Bertsy Goic, DrawInScience

Cellular navigation is further complicated by the structural complexity of tissues, where cells must detect and interpret chemical gradients at microscopic scales, while adapting to mechanical constraints, and avoiding physical obstacles. Failed or inefficient migration has serious consequences, as seen in cancer, where immune cells often struggle to penetrate dense tumour microenvironments. While the biochemistry of gradient sensing is increasingly well understood [12], and theoretical models such as run-and-tumble particles (RTP), active Brownian particles (ABPs), and active Ornstein-Uhlenbeck particles (AOUP) have provided useful frameworks for active motion [13–15], it remains unclear how these models map onto real immune cell behaviour in biologically relevant complex environments.

Here, we combine theory and experiment to investigate how neutrophils and DCs navigate in complex environments during chemotaxis. Using *in vitro* collagen matrices with controlled chemokine gradients, we track single-cell trajectories over extended time and length scales. This integrated multiscale approach allows us to dissect how physical constraints, gradient properties, and cell-intrinsic programs collectively shape migration from cell to tissue scale. By directly comparing two functionally distinct immune cell types under physical conditions, we uncover general and cell-specific strategies for efficient chemotaxis in tissue-like environments.

## Distinct reorientation dynamics underlie immune cell chemotaxis

To mimic the tissue environment, we used collagen gels, the major component of interstitial tissues, at densities that impose different confinement levels on cells, as observed in tissues. For this, we let a mixture of cells and collagen gel polymerize in a microfabricated chip as described previously [16], see Fig. 2a and methods section for details. In this set-up, both DCs and neutrophils show a clear bias towards the chemokine source (Fig. 2b and c). As the two cell types differ in speed and size, different collagen concentrations were chosen to ensure that neutrophils and DCs cover a comparable distance up the gradient (Fig. 2d and e).

**Fig. 2.**
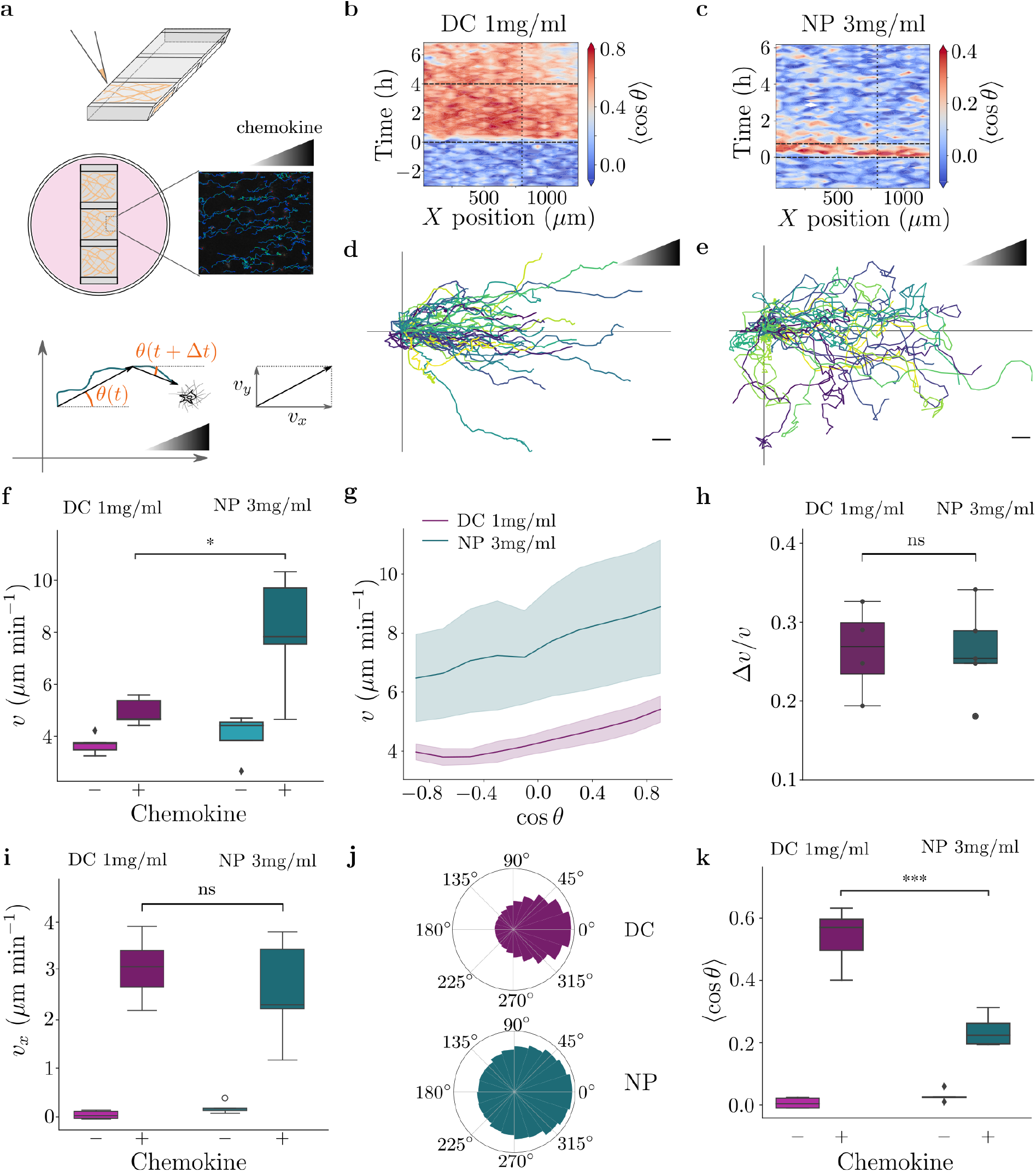
Chemotaxis of dendritic cells (DC) and neutrophils (NP) in porous environments. **a**, Cells embedded in a collagen gel are loaded in a microfabricated chip, which is surrounded by medium (with chemokine and drugs where necessary). The orientation *θ* is defined with respect to the orientation of the chemokine gradient, which is aligned with the positive *x* axis. **b**, Kymograph of the directionality, ⟨ cos(*θ*) ⟩, of DCs in 1 mg/ml collagen gel for a representative experiment. Chemokine addition is indicated by the dashed horizontal line at time 0 h. The trajectories are included up to the second horizontal line. The analysis is restricted to the region closest to the chemokine source, indicated by the dotted vertical line and right edge. **c**, Kymograph of the directionality, ⟨cos *θ* ⟩, of neutrophils in 3mg/ml collagen gel for a representative experiment. **d-e**, Example trajectories of DCs and neutrophils migrating in 1mg/ml and 3mg/ml collagen gels, respectively. The scale bar corresponds to 20 µm. **f-g**, Instantaneous speed (*v*), and instantaneous speed as a function of orientation, cos *θ*. **h**, Relative speed change, **i**, Instantaneous speed along the gradient direction, *v*_*x*_, **j**, Distribution of orientations *θ*, **k**, Average directionality, ⟨cos *θ*⟩. N=5 for neutrophils and DCs.

Consistent with previous studies [17, 18], we find that neutrophils significantly increase the mean magnitude of the instantaneous velocity, *v*, in the presence of chemokine (Fig. 2f). While the magnitude of this increase is smaller for DCs, the relative increase of speed magnitude is comparable for both cell types when they are oriented towards the chemokine (Fig. 2g and h), where *θ* = 0 is the orientation of the gradient along the x axis. In principle, such a bias in orientation-dependent speed, *v*(*θ*), is sufficient to induce a chemotactic response, even if the orientation *θ* remains isotropic (see SI). Although the magnitude of instantaneous velocity is significantly smaller for DCs than for neutrophils, the magnitude of the velocity component along the gradient direction, *v*_*x*_ = ⟨ *v*(*θ*) cos *θ* ⟩, does not differ between the cell types. Thus, the chemotactic efficiency of DCs must be improved by other means. Indeed, in addition to a bias in the speed, both cell types also show a bias in the distribution of orientations, *P* (*θ*) (Fig. 2j), and the average directionality, ⟨cos *θ*⟩, (Fig. 2k). A skewed distribution *P* (*θ*) is in principle also sufficient to independently induce a chemotactic response. Yet, observing such a bias in the distribution of orientations does not explain how the cells navigate locally to control their orientation nor is it sufficient to conclude, e.g., that cells actively turn towards a gradient, or adjust the magnitude of angular fluctuations.

To understand how the cells achieve this bias in orientation, we examine the change in orientation as a function of the current orientation of the cell, i.e. whether the cell adjusts the angular velocity, *ω* = *∂θ/∂t* in response to the orientation *θ*. Analysis of trajectories during the initial random migration period shows that the collagen gels are isotropic and do not bias the migration (Fig. 3a-b). When the chemokine is introduced, the conditional probability density of the angular velocity shows a striking difference between neutrophils and DCs. While there is only a slight bias in the average angular velocity for neutrophils, DCs show a clear linear dependence on the orientation, which implies an effective torque that acts to reorient the cell towards the chemokine (Fig. 3c, d and e). In addition to the median of the angular velocity, we found that the cells also modulate the fluctuations of the angular velocity depending on the local orientation, which can be quantified by the standard deviation, *σ*_*ω*_(*θ*). When either neutrophils or DCs are oriented towards the gradient, the magnitude of fluctuations is significantly reduced compared to the random migration. In fact, the minimum of *σ*_*ω*_(*θ*) coincides with *θ* = 0, i.e. when the cell is aligned with the orientation of the chemokine gradient. However, while neutrophils moving away from the gradient, i.e. *π/*2 *<* |*θ*| *< π*, show an angular noise similar to the random case, DCs display an increase in the magnitude of angular fluctuations for these angles.; that is, DCs turn more when moving away from the gradient than during random migration.

**Fig. 3.**
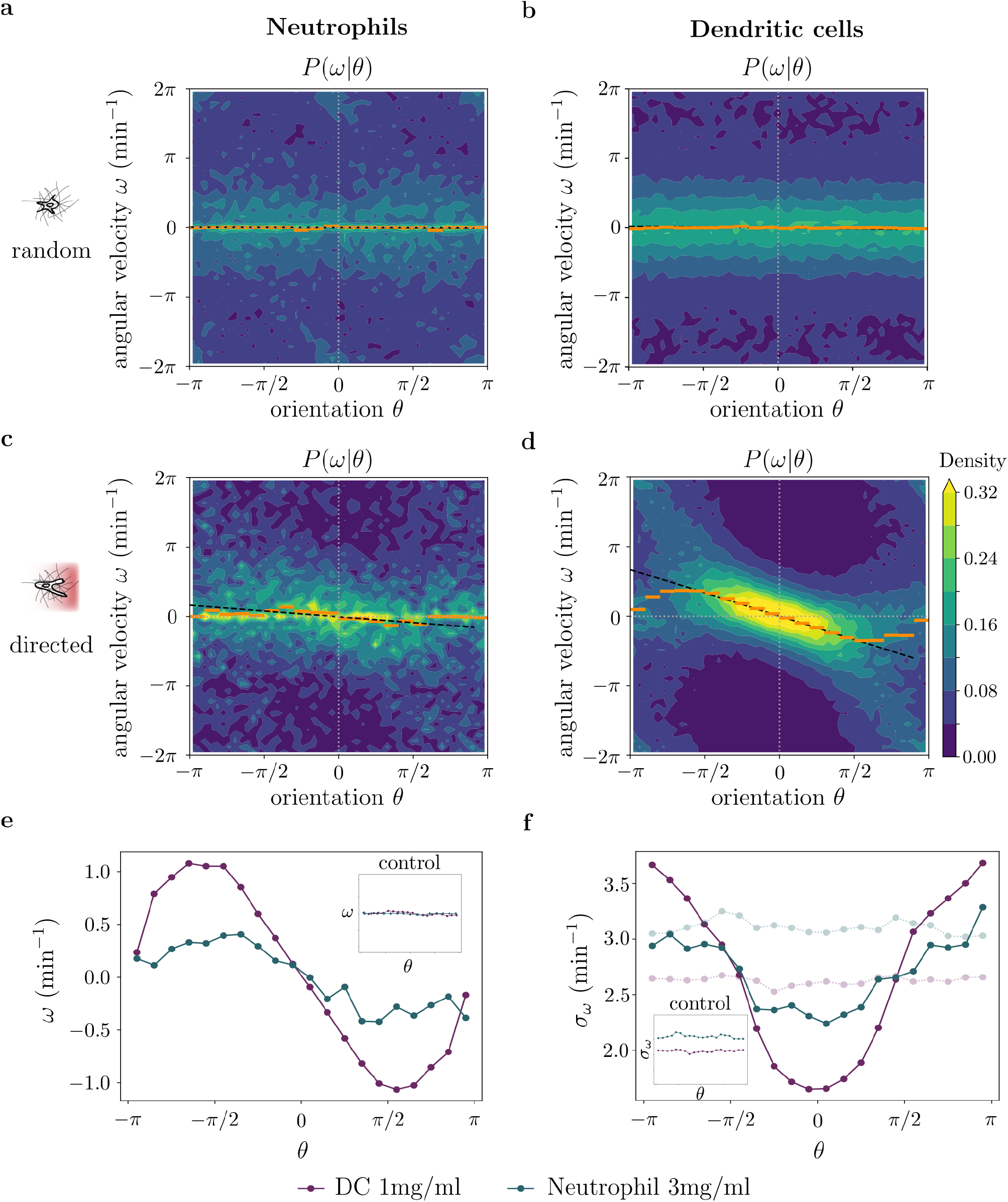
Torque and coloured noise govern orientation dynamics. Conditional probability distributions of angular velocity *ω* given the orientation *θ* for a representative experiment. **a–b**, Neutrophils and DCs in the absence of a chemokine gradient. **c-d**, Same cell types in the presence of a chemokine gradient. Orange lines, binned medians; dashed black lines, linear fits. **e**, Median *ω*(*θ*) in a gradient (inset: random migration). **f**, Standard deviation *σ*_*ω*_(*θ*) in a gradient (inset: random migration). Neutrophils: n=1,163 (random), n=676 (directed); DCs: n=1,593 (random), n=2,659 (directed).

A reason for these differences could stem from the fact that we used different concentrations of the collagen gels for the two cell types. We thus performed the same experiments with DCs in a denser collagen gel, at the same concentration used for neutrophils. Although the denser collagen gel slowed down the DCs, their capacity to chemotact and the metrics of the angular velocity were not changed (Fig. 4 and Supp. Fig. 4). The biased speed increase is therefore dispensable to the chemotactic strategy of DCs since they maintained their ability to actively control their orientation despite the increased complexity of the environment. This observation confirms that DCs and neutrophils clearly show different dynamics in their control of orientations, which do not depend on collagen density (Supp. Fig. 4).

**Fig. 4.**
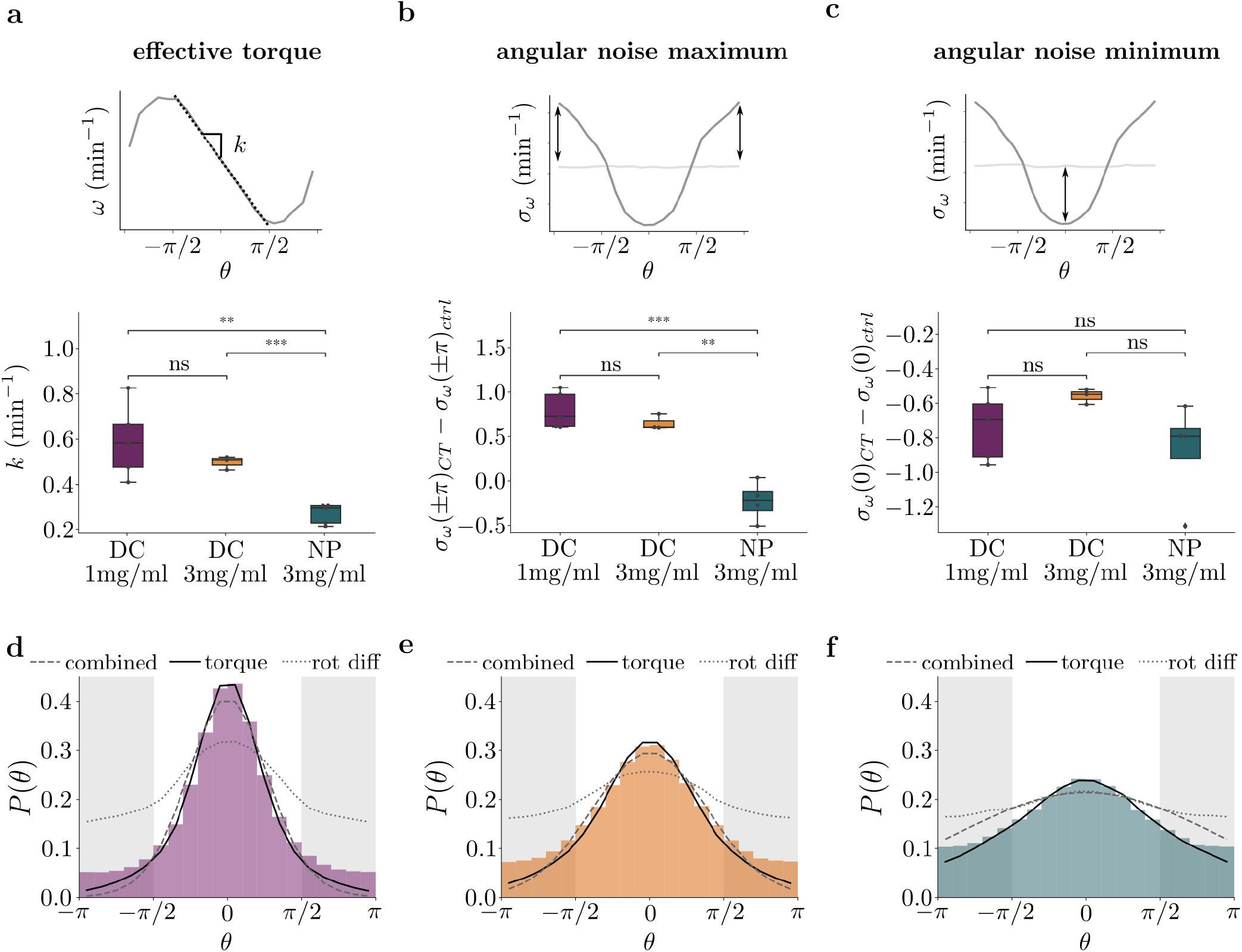
Deterministic and stochastic chemotaxis strategies. **a**, Effective torque parameter *k* **b** Difference between the angular noise in a chemokine gradient and during random migration in the direction opposite of the chemokine gradient **c**, Difference between the angular noise in a chemokine gradient and during random migration in the direction of the chemokine gradient. N=5 for neutrophils and DC 1mg/ml, N=3 for DC 3mg/ml **d-f**, Experimentally obtained distribution of orientation *P* (*θ*) compared to theoretical predictions (lines)

## Deterministic and stochastic orientation control define distinct chemotactic modes

The observed dependence of the angular velocity on orientation suggests that the cells employ two components to adjust the orientation: a deterministic and a stochastic part. In the context of stochastic processes, both deterministic and stochastic forces can in principle independently lead to a bias in the steady state distribution of orientations, but it is unclear which process is more important in determining the chemotactic response of the cells. To identify their relative importance, we turn to theoretical arguments. For a generic (Markovian) stochastic process, the time evolution of the probability density function of an observable such as the orientation can be described on general grounds by a Fokker-Planck equation. In its general form, it can be written as

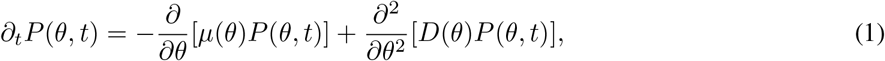

where *µ*(*θ*) and *D*(*θ*) are the drift and (biased) diffusion term, respectively. The drift arises from a deterministic bias in the angular velocity, while the biased diffusion term stems from a bias in the amplitude of fluctuations. Thus, both drift and diffusion terms can be inferred directly from the experimental mean and standard deviation of the angular velocity in Fig. 3, according to *µ*(*θ*) = ⟨Δ*θ/*Δ*t*⟩ = ⟨*ω*(*θ*)⟩ and 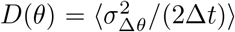. For analytical purposes, we approximate the drift term according to the linear dependence identified in Fig. 3e as *µ*(*θ*) = −*kθ*, which introduces *k* as an effective torque parameter. Solving the FPE gives the stationary distribution of the orientation, *P* (*θ*), as

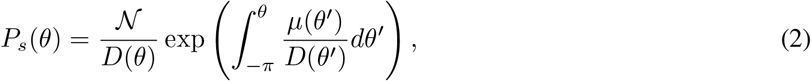

where 𝒩 is a normalization constant. We can compare this theoretical prediction to the experimentally obtained distribution of orientations. Considering the stochastic only (i.e. *µ*(*θ*) = 0 while *D*(*θ*)) and the deterministic only case (i.e. *D*(*θ*) = *D* while *µ*(*θ*)) we ask whether they are sufficient to explain the observed distribution. As the linear relationship in Fig. 3, which implies an effective torque, breaks down outside the range [−*π/*2, *π/*2], we determine 𝒩 not directly by normalizing the expected probability over the entire range [−*π, π*]. Instead, we determine it by fitting Eq. (2) to the experimentally determined *P*_*s*_(*θ*) restricted to the range [−*π/*2, *π/*2], with 𝒩 being the only fitting parameter.

For DCs, we find that the deterministic part is sufficient to explain the experimentally observed distribution of orientations in dense collagen gels (Fig. 4). Conversely, for the neutrophils, neither the deterministic nor the stochastic part are on their own sufficient to explain the distribution of orientations, but the combined mode shows a good agreement with the experimental data. In conclusion, the model supports the hypothesis that neutrophils and DCs are using different chemotactic strategies to bias their orientation, with DCs employing mainly an active torque to steer their direction of migration along the chemotactic gradient, while neutrophils also make use of a modulation of the amplitude of fluctuations of their angular velocity. It is however unclear what are the main factors – either from internal cellular, or external environmental constraints – that control the deterministic and stochastic part of the cell response.

## Chemotactic strategy depends on environmental structure and cytoskeletal organisation

Since increasing the complexity of the environment by increasing the density of collagen gel did not significantly reduce the capacity of DCs to efficiently chemotact using an effective torque (Fig. 4a), we wondered whether the torque is maximized in the absence of obstacles. As DCs require confinement to migrate efficiently, we used a quasi 2D confined migration assay with a roof lower than the cell diameter in which we establish a chemokine gradient (Fig. 5a & methods for details). In this environment, DCs displayed robust directed migration (Fig. 5b&c). In the absence of gels, DCs strongly increased their speed when they were aligned with the gradient (Supp. Fig. 5) showing that the biased speed increase is maximized in the absence of obstacles. However, despite the clear evidence of chemotaxis, the effective torque was surprisingly strongly reduced in DCs migrating in the confined 2D environment to levels comparable to neutrophils in collagen gel (Fig. 5d). Thus, DCs significantly altered their chemotactic strategy in response to the environment.

**Fig. 5.**
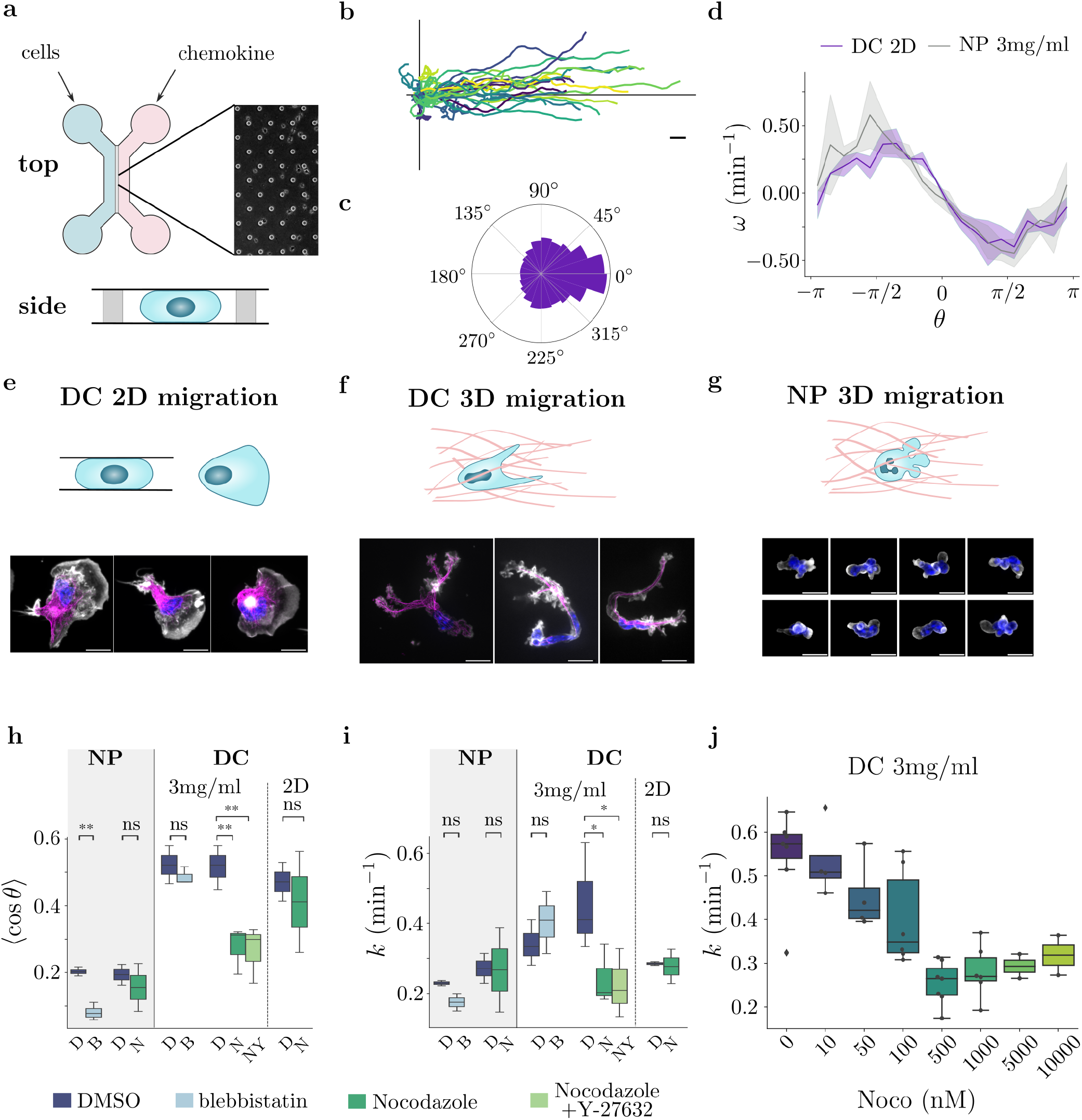
Chemotactic torque depends on environment and microtubules. **a**, 2D migration setup; **b**, Example DC trajectories in a CCL19 gradient in 2D; scale bar 20 µm; **c** Distribution of orientations; **d**, Angular velocity for DCs in 2D and neutrophils in 3D collagen (N=5 for neutrophils, N=2 for DCs in 2D); **e-g**, Representative cell morphologies in 2D DCs, 3D DCs, and 3D neutrophils; scale bar, 10 µm; grey scale: actin; blue: Hoechst; magenta: microtubules. **h-i**, Effects of environment and drug treatments on directionality and effective torque parameter. D: DMSO, B: blebbistatin, N: Nocodazole, NY: Nocodazole + Y-27632 (neutrophils: N=2 for all conditions; DCs: N=2 for 2D experiments, N=3 for all other conditions). **j**, Dose response of effective torque parameter to nocodazole (DCs in 3mg/ml collagen gel). N=3 for 0-1 µM and N=2 for 5 µM-10 µM.

We next turned to identify the cellular mechanisms that facilitate this adaptation in response to the complexity of the environment. A clear difference between DCs in collagen gel and 2D confined environments is the cell morphology. In collagen gel, DCs take on a characteristic branched morphology (Fig. 5f). DCs extend multiple long branches during migration, and in order to move forward, one of these branches has to be stabilised while the others are retracted [19]. In the 2D environment, on the other hand, DCs adopt a large and continuous travelling front, consistent with previous reports [20,21]. As a consequence of this different shape, the cytoskeleton, including the actomyosin as well as the microtubule network, displays a very different organisation. In 2D we observed a fan-like MT network, where the microtubule tips hardly reached the actin cortex at the travelling front (Fig. 5), whereas extending branches in collagen gels were populated by bundles of microtubules [22]. Since microtubules have previously been shown to control the decision between competing branches in DCs [23], we hypothesized that the differences in the microtubule architecture could explain the loss of torque observed in 2D confined environment, as well as the absence of a strong torque effect in neutrophils, which never display large microtubule bundles.

Perturbing microtubules polymerization using Nocodazole showed a clear decrease in the directionality for DCs migrating in a collagen gel, and reduced the torque effect in a dose dependent way, reaching levels observed for neutrophils (Fig. 5), and DCs in 2D confined environments. On the other hand, Nocodazole had no/little effect on DC chemotaxis in 2D confined environments, which supports the idea that it specifically disturbs the torque effect. As a decrease in the amount of microtubules also leads to an activation of the RhoA exchange factor GEF-H1 [24], nocodazole also increases the cell contractility, altering the cell shape and increasing speed [25]. To avoid this additional effect, we treated cells with the Rho kinase inhibitor Y-27632 in addition to nocodazole. In this condition, cells kept their shape and speed, but the torque effect was not recovered (Fig. 5i). This suggests that microtubules play a specific role in the effective torque displayed by DCs in collagen gels, which is independent of their role in contractility. In addition to the decrease in the magnitude of the torque, the bias in the angular noise is also reduced upon Nocodazole treatment, in particular the increase when cells move away from the gradient, which is specific to DCs (Supp. Fig. 5). Overall Nocodazole treatment shifts the chemotactic response of DCs in collagen to the response employed by neutrophils.

Contrary to DCs, the chemotactic response of neutrophils is not affected by Nocodazole treatment. Neutrophils have high levels of contractility, resulting in a more rounded cell morphology without long branches containing microtubules. Perturbations that affect the actomyosin network, e.g. blebbistatin, reduce the ability of cells to migrate and lead to a reduction in speed. Consequently, the directional speed increase in neutrophils is lost (Supp Fig. 5). Furthermore, blebbistatin reduced the directionality but not the effective torque effect for neutrophils, indicating an effect on the directional noise instead. Conversely, DCs still showed a clear directional response, as the effective torque, which dominates their chemotactic capacity, was hardly affected by these treatments decreasing contractility.

Put together, the two specific elements of DC chemotaxis, the effective torque, and the increase in directional noise when moving away from the gradient, both depend on microtubules. On the other hand, the decrease in directional noise for cells migrating towards the chemokine, which dominates the chemotactic capacity of neutrophils, depends on acto-myosin contractility. As a consequence, the capacity of DCs and neutrophils to display directed migration is affected by different drugs, targeting different cytoskeletal structures.

We can summarise the effect of different treatments and conditions in a phase space spanned by the stochastic and deterministic part of the response. To quantify the decrease in angular noise, the experimentally observed *D*_*r*_(*θ*) can be approximated by

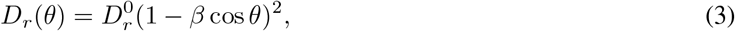

which introduces the phenomenological parameter *β* that controls the relative amplitude of noise modulation. In addition, the theoretically predicted directionality can help to assess the relative importance of the torque and angular modulation in the chemotactic strategy. For this, the directionality 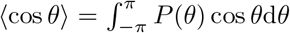 is derived using *P* (*θ*) given by Eq. 2 for a chemotactic strategy based exclusively on a torque effect or on biased angular fluctuations (see SI).

To compare the different experimental conditions to the theory, the torque parameter has to be non-dimensionalized by the baseline angular noise, 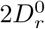 (see SI for details), introducing the normalized torque 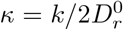. The phase space defined by the two parameters that quantify the normalized effective torque and decrease in noise - *κ* and *β*, respectively - is shown in Fig. 6. The relative strength of the directionalities predicted by the different strategies, Ψ, is shown as shading, with equal directionality (i.e. Ψ = 0.5) shown as a solid black line. If a condition is on the left-hand-side of the line, the parameters obtained from experiments are such that the biased angular noise is predicted to have a stronger impact in the chemotactic strategy.

**Fig. 6.**
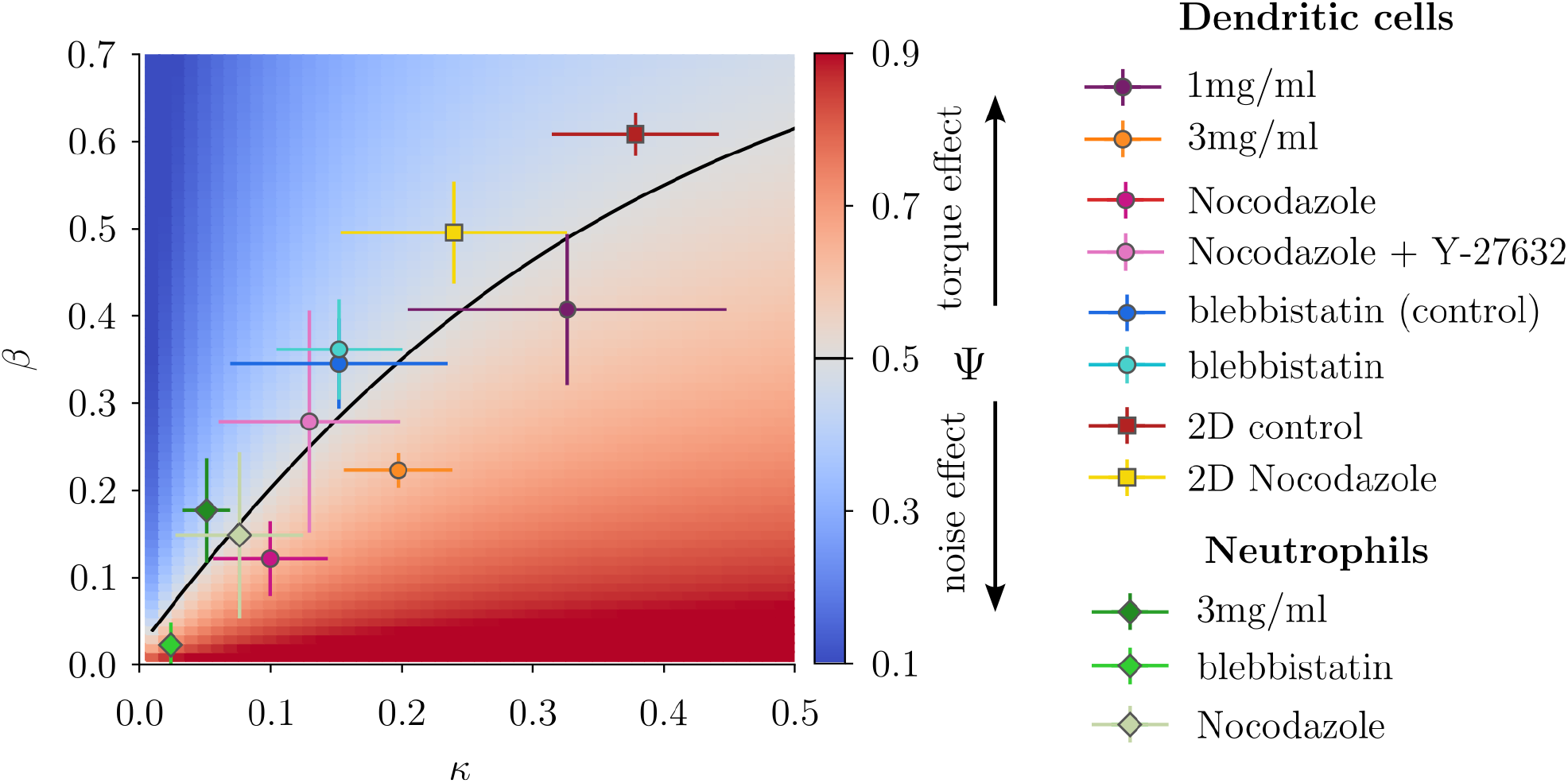
Chemotactic phase space. Phase diagram of chemotactic strategies in the space defined by the rotational diffusion bias, *β*, and normalized torque strength, *κ* = *k/*2*D*_*r*_. Data points represent mean with standard deviation. Shading indicates the ratio Ψ between the drift predicted by the torque-only strategy and the sum of drifts from torque and rotational diffusion (see SI for details). The ratio thus shows which strategy would lead to a stronger effect if it was chosen exclusively. The solid contour marks equal strength of the chemotactic response predicted by the two different mechanisms (Ψ = 0.5).

The phase space in Fig. 6 combines many experimental conditions, including different environments and pharmacological perturbations. Yet, the data points roughly follow the black line of equal strengh of the chemotactic strategies. While they all stay close to this transition, they fall on different sides. Neutrophils fall on the left-handside, indicating that angular noise modulation has a slightly stronger effect. Conversely, DCs in loose and dense collagen gel are situated on the right-hand-side of the transition as the torque has a stronger effect. Even though they remain on this side, treatment with Nocodazole clearly shifts DC behaviour closer to neutrophils in the phase space. The larger effect of drugs affecting contractility on neutrophils than on DCs is also clearly illustrated by this graph. Nevertheless, the positive correlation between *β* and *κ* predicted by Fig. 6 indicates that the underlying cellular processes are related.

Strikingly, DC migration in 2D shows the largest normalized torque effect, *κ*, while in Fig. 5 the torque parameter, *k*, was shown to be small. The difference is due to the normalization as DCs migrating in 2D show a large persistence, and, thus, small 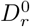 (see Supp. Fig. 6 for plot without normalization). Furthermore, perturbing microtubules affects the persistence of cells migrating in 2D, but not the chemotactic torque parameter, *k*, which explains the apparent effect of Nocodazole treatment in Fig. 6. While DCs migrating in 2D show a larger normalized *κ*, they are situated on the LHS of the black line and their behaviour is still predicted to dependent more strongly on the angular noise modulation than on the torque effect - unlike DCs migrating in collagen gel. Indeed, in collagen gel, cells are faced with frequent reorientations imposed by the complex structure of their environment, which stronger chemotactic torque can counterbalance.

## Population-level consequences of deterministic and stochastic chemotactic strategies

We have shown that, although they both have critical functions in the immune response, neutrophils and DCs employ different cytoskeletal components to achieve different chemotactic strategies in complex 3D environments. Yet, it remains unclear why such closely related cells would have evolved to use different strategies for a seemingly similar goal. In order to understand the benefit of these different strategies, we introduce Langevin equations to model single-cell trajectories. Taking into account the orientation dynamics defined by the system described in Eq. (1), we write

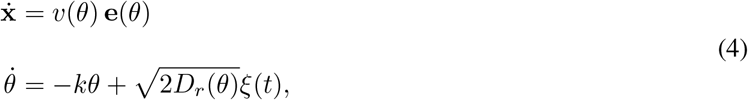

where dot denotes the time derivative, we introduce the unit vector **e** = [cos *θ*, sin *θ*], and the white noise in Eq. (4) obeys ⟨*ξ*_*r*_(*t*)⟩ = 0 and ⟨*ξ*_*r*_(*t*)*ξ*_*r*_(*t*^′^) ⟩ = *δ*(*t* − *t*^′^). Note that the equivalence with the Fokker Plank equation Eq. (1), inferred from data, imposes here the Ito prescription without ambiguity. A pure stochastic strategy then consists of 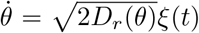. On the other hand, a cell that uses exclusively a deterministic strategy adjusts the orientation via 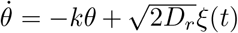 with the effective torque parameter *k*. Because we focus on the orientational dynamics, we will assume *v*(*θ*) = *v* constant hereafter (see SI for effect of speed modulation).

From the single-cell model it becomes clear that the two different strategies impose different requirements on the cells ability to resolve a gradient. The bias in the noise, *D*_*r*_(*θ*), needs to be symmetric around the target direction *θ* = 0 to be effective. Cells therefore only need to be able to resolve the chemokine gradient along cos *θ*, i.e. only a projection of the gradient instead of the full gradient. This projection can be along the major cell axis or, if the cells are not large enough, along a constant travel direction as was observed for bacteria such as *E*.*coli* (Fig. 7a). The deterministic torque strategy, however, requires that the cells are able to resolve the entire gradient direction, i.e not just along the polar axis but also the lateral axis, in order to turn directly towards the gradient. It thus requires a refined sensing mechanism, and typically a larger spatial extension beyond the cell polarity axis. Both strategies introduce a bias in the orientation, which leads to a biased displacement via 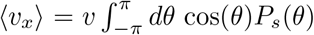 Notably, both the deterministic and stochastic strategies can lead to the same drift strength with an appropriate choice of parameters (Fig. 7).

**Fig. 7.**
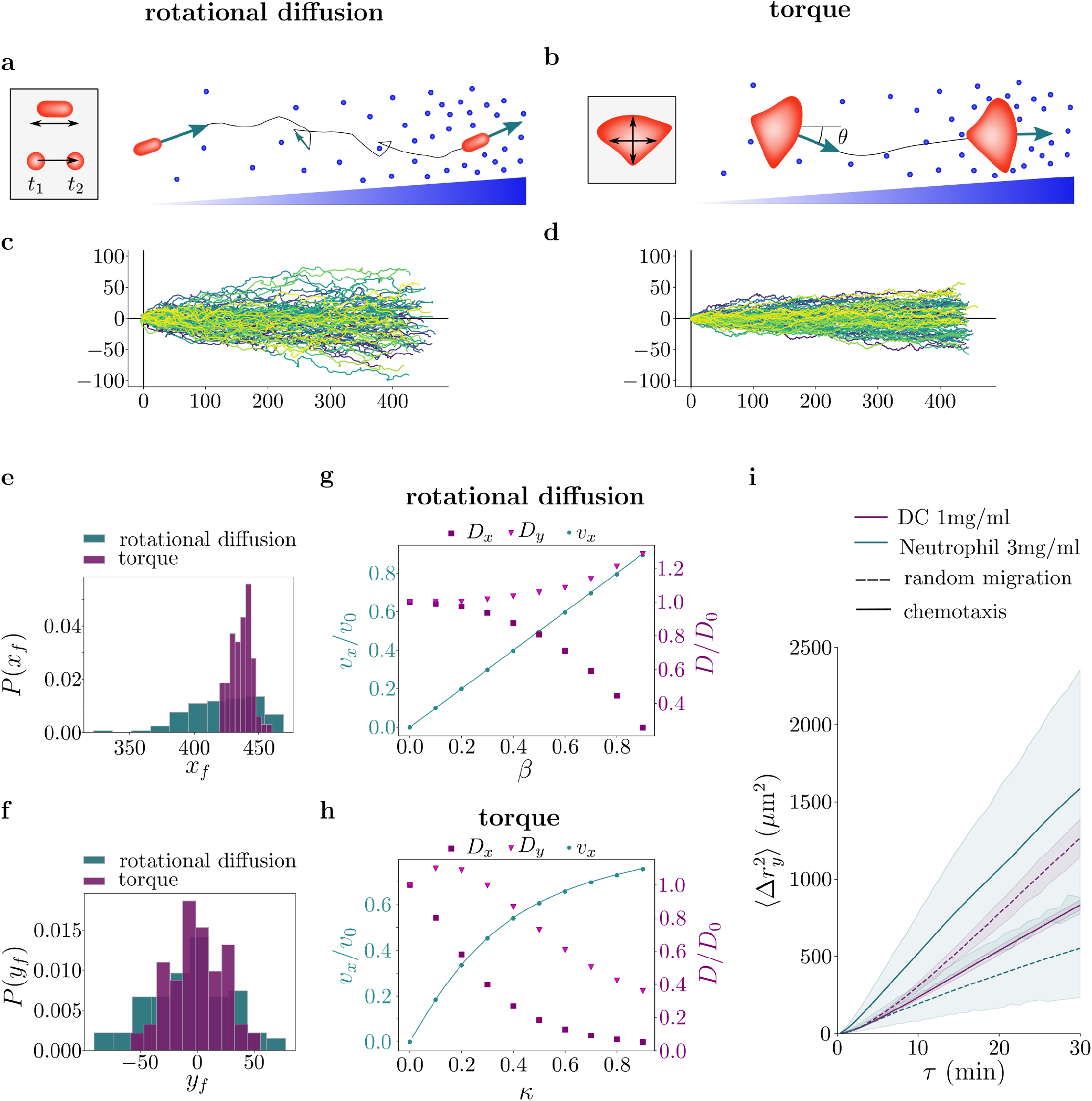
Population-level consequences of deterministic and stochastic chemotaxis strategies. **a**, Schematics of chemotactic response based on cos *θ*, which is determined either by direct sensing along the length of a polar particle or approximated by memory of measurements over a time interval [*t*_1_, *t*_2_] **b**, biasing the orientation towards a gradient *θ*, which is measured directly along the axes of an extended particle. **c**, Simulated trajectories for a polar particle with *D*_*r*_(*θ*) = (1 − *β* cos *θ*)^2^, where *β* = 0.7. **d**, and a torque particle with *κ* = 0.8. The choice of parameters ensures equal chemotactic drift strength for both strategies. **e**–**f**, Final position distributions in the *x* (gradient) and *y* (perpendicular) directions. **g**-**h**, Drift velocity *v*_*x*_*/v*_0_ and diffusion coefficients *D*_*x*_, *D*_*y*_ as a function of the biasing parameter for both strategies (symbols, simulations; lines, analytical results). **i**, Meansquared displacement perpendicular to the gradient, with and without chemokine gradients (solid and dashed lines).

If a cell is able to resolve the full gradient, it might be tempting to assume that it would thus use a deterministic strategy. However, the two-point dynamics of the two processes are not equivalent, even if they can lead to similar distributions *P* (*θ*). A reasonable approximation of the experimentally observed bias in the rotational diffusion coefficient is the *D*_*r*_(*θ*) introduced in Eq. (3). Increasing the bias by increasing *β*, increases the drift in ⟨*x*⟩, but at the same time increases the spread of the population prependicular to the gradient direction (Fig. 7g). Conversely, if the angle is biased as a Ornstein-Uhlenbeck process, 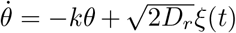, increasing the drift in *x* by increasing *k* (while *D*_*r*_ is assumed to be constant) leads to a decrease in the spread of the population both perpendicular as well parallel to the gradient direction (Fig. 7h). This means that, contrary to polar particles, cells able to display an effective torque strategy can move up gradients without showing a dispersion in the axis perpendicular to the gradient and increasing their bias will even further decrease their dispersion. Furthermore, the spread along the gradient direction is strongly reduced for particles using a torque effect, whereas it decreases to a smaller extent for particles using a biased noise term. This can, for example, lead to differences in reaching a target distance, *x*_*f*_, along the gradient direction (Fig. 7e). In conclusion, both chemotactic strategies give rise to a bias in the distribution of orientations and, thereby, a chemotactic drift, but display marked differences in the population spread. Thus, cells might choose one or the other based on the population behaviour that is more suitable.

A prediction of the model is then that the spread in the direction perpendicular to the chemokine gradient should differ between the two cell types as the chemotactic response of DCs is dominated by the deterministic response, whereas the neutrophil response is strongly influenced by the stochastic response. This theoretical prediction is confirmed by the experimental data, where we find that the mean squared displacement in the perpendicular direction is reduced in the chemotactic case compared to the random case for the DCs (Fig. 7i). Neutrophils, on the other hand, spread more in the perpendicular direction during chemotaxis than during random migration.

## Discussion

Despite both being rapidly migrating immune cells, we found that dendritic cells and neutrophils exhibit distinct chemotactic strategies. Specifically, we observed that DCs display a clear effective torque, whereas neutrophils predominantly exhibit an angle-dependent modulation in speed and angular noise. This torque is lost in DCs treated with nocodazole or when migrating in free space devoid of obstacles (2D confinement). The likely origin of the torque lies in the branched morphology of DCs and their interaction with the surrounding matrix.

Microtubules may contribute to the torque by stabilizing specific cellular protrusions and influencing the direction of new ones. Imaging showed DCs extending multiple long protrusions at the leading edge. As the cell advances, all but one protrusion retracts, and the stabilized one sets the new direction. This directional choice is biased toward protrusions aligned with the chemokine gradient, a bias that diminishes upon microtubule disruption (Supp. Fig. 7). When treated with nocodazole and Y-2763, DCs form more protrusions pointing away from the chemokine source, and selection of the stabilized branch becomes more random, increasing angular noise. These findings suggest that microtubules influence both the initiation and stabilization of protrusions, consistent with their established role in cell polarity and long-range intracellular transport [26]. Additionally, microtubule-associated regulators like RhoA GEF-H1 may also play a role [27]. Notably, the effect of nocodazole saturated at low doses, where microtubules are still present, but the dynamics of growing microtubule ends are strongly impaired [28, 29]. Microtubules may hence play a role in the effective torque by interacting with the distal growing tips of cellular protrusions, where microtubule + ends are enriched. Several molecular mechanisms have been proposed for such interactions [30–33], and could be candidates for the mechanisms underlying the effective torque.

In contrast, neutrophil chemotaxis is driven by increased speed and biased noise in response to a chemokine gradient, and microtubules have little impact on the chemotactic capacity of neutrophils. Consistently, neutrophils do not extend long protrusions, and their microtubule cytoskeleton appears less developed, being confined to the cell center. Instead, perturbing contractility strongly impacts not only the speed but also the biased angular noise. The role of Myosin II in directed migration is well established in neutrophils [34, 35], as well as in other cell types [36], suggesting that other cell types also use a comparable strategy rather than torque based navigation. Because the effective torque was insensitive to contractility inhibition, such treatments have little effect on DC chemotaxis.

The relative contribution of the identified chemotactic strategies can be quantified by extracting two parameters from single-cell trajectories – one measuring the bias in the orientational noise and the other measuring the linear dependence of the rotational speed on the direction of migration, defining the strength of an effective torque. This provides a generic metric for comparing cell types, drug treatments, and the impact of the migration environment, complementing classical metrics like angular bias, speed, and persistence, which are insufficient to distinguish chemotactic strategies. Using this metric, our study highlights that distinct cytoskeletal components support different chemotactic strategies.

In addition to different cytoskeletal components, the different chemotactic strategies also pose different requirements on sensing and gradient resolution. The torque requires spatial detection of the full gradient, implicating a more spatially extended cell shape, and a mechanism for comparing signals from different protrusions. In collagen, DCs indeed display a more elongated and branched cell shape, consistent with the requirements for a torque strategy, while neutrophils have a round cell shape. On the other hand, an estimate of a projection of the gradient is sufficient for a directional increase in speed as well as a reduction in orientational noise. The projection of the gradient can be estimated either along the cell polarity axis, or as a temporal comparison along the migration path. While the temporal gradient estimation is generally considered restricted to prokaryotes [37], it would be sufficient to achieve the chemotactic strategy of neutrophils [38, 39]. However, even if cells are able to perform a full estimation of the gradient, they might still choose a polar strategy such as the biased angular noise due to the macroscopic consequences of the different chemotactic strategies. Using active particle models, we showed that a bias in noise leads to a larger spread of the population than the torque strategy. Depending on the environment, differences in, e.g., the mean arrival time at a target can arise. Thus, the two strategies are not simply interchangeable.

Altogether, our findings suggest that directed cell motility is adapted to the cell’s function, beyond mere limitations of size or shape. Neutrophils, when dealing with infected tissues, respond to heterogeneous, time-varying chemical gradients. A chemotactic strategy that leads to an increased spread could ensure that some cells reach the target while others remain receptive to other potential areas of infection. At the same time, the response to the original threat remains effective as neutrophils arrive in large numbers. In contrast, only very few DCs patrol tissues. Once they have encountered a danger signal and matured, they respond to chemical gradients that lead them to a lymphatic vessel. They then reach a lymph node, where they trigger the adaptive immune response by presenting the signal to T cells. It is thus crucial that most DCs that have picked up a danger signal reach the nearest lymphatic vessel — a fixed spatial target — as quickly as possible. An effective torque is an appropriate strategy for this purpose.

## Supporting information

Supplementary model information

## Author contributions

T.J. performed most DC experiments, analysis and theoretical modelling; M.D. performed neutrophil experiments; A.M. and P.J.S. performed initial DC experiments; M. B. helped with DC experiments; L.W. performed photolithography; L.B. helped with image analysis; P.J.S. performed immunostaining on DCs; P.V. designed the experimental approach to study leukocyte chemotaxis in 3D; T.J., R.V., and M.P., designed the study and wrote the manuscript. All authors contributed to discussions and the final version of the manuscript.

## Acknowledgements

We thank Ana-Maria Lennon Dumenil (Institut Curie, Paris, France) for helpful discussions and sharing mice stocks, Bianca Cali (Institute of Oncology Research, Bellinzona, Switzerland) for initial help in setting the experimental system to study the directed migration of neutrophils in collagen gels, and Bertsy Goic, DrawInScience, for preparing the illustration used in Fig. 1. This work benefited from the technical contribution of the joint service unit IPGG Technological Platform (CNRS UAR 3750).

## Methods

### Mice

Male and female C57BL/6J (B6) mice aged 8–16 weeks were used. Animals were either wild-type (C57BL/6J inbred strain; Charles River) or transgenic lines expressing Lifeact–EGFP (kind gift from M. Sixt) or Myosin IIA–GFP (kind gift from R. S. Adelstein). None of these strains exhibited any deleterious phenotype; therefore, their breeding and maintenance were not subject to project evaluation. Mice were housed under specific pathogen–free conditions in the animal facility of Institut Curie, in accordance with European and French regulations for the protection of vertebrate animals used for scientific purposes (Directive 2010/63; French Decree 2013-118). Animals were euthanized by cervical dislocation performed by trained and authorized personnel.

### Bone-marrow derived dendritic cell cultures

DCs were obtained as described previously [40]. In brief, bone marrow cells were isolated by flushing both whole legs from 6-to 8-week-old mice and cultured for 10 days in IMDM (Sigma-Aldrich) supplemented with 10% decomplemented and filtered fetal bovine serum (FBS; Biowest), 20 mM L-glutamine (Gibco), 100 U m1^−1^ penicillin–streptomycin (Gibco), 50 µM *β*-mercaptoethanol (Gibco), and 50 ng m1^−1^ granulocyte–macrophage colony-stimulating factor (GM-CSF)-containing supernatant obtained from transfected J558 cells, as previously described [41]. On days 4 and 7, cells were detached using 5 mM PBS–EDTA and replated at 0.5×10^6^ m1^−1^. At day 10, semi-adherent cells were collected by gentle flushing after removal of non-adherent cells. For stimulation, BMDCs were treated on day 10 with 100 ng m1^−1^ Lipopolysaccharide (Salmonella enterica serotype Typhimurium; Sigma) for 30 min, washed three times with complete medium, replated in fresh medium, incubated overnight, and used on day 11.

### Mouse neutrophil isolation and culture

Neutrophils were obtained as previously described [42]. Briefly, single-cell suspensions were prepared from bone marrow flushed from the femurs and tibias of mice. Neutrophils were isolated by immunomagnetic negative selection using the MojoSort Neutrophil Isolation Kit (BioLegend, #480058) according to the manufacturer’s instructions. After isolation, cells were cultured overnight at a concentration of 1 × 10^6^ cells m1^−1^ in RPMI 1640 medium supplemented with 10% fetal bovine serum (FBS), 100 U m1^−1^ penicillin–streptomycin, and 50 ng m1^−1^ recombinant granulocyte–macrophage colony-stimulating factor (GM-CSF; PeproTech, #315-03-20UG).

### Drug treatments

The following pharmacological inhibitors and chemical compounds were used: Nocodazole (Sigma-Aldrich, #M1404); blebbistatin (Abcam, #ab120425); Y-27632 (Biotechne, #1254/10). Nocodazole was used at 0.5 µM (DCs) and 10 µM (neutrophils) in Fig.5h-i. Blebbistatin was used at 10 µM (DCs) and 50 µM (neutrophils). Y-27632 was used at 1 µM (DCs and neutrophils). Medium was supplemented with DMSO (Sigma-Aldrich, #D2438) at the appropriate concentration in control experiments when DMSO was used as a solvent.

### Photolithography and PDMS chip preparation

All microdevices used in this study were designed and fabricated as follows. Chrome photomasks were produced by JD Photo Data (UK) based on custom designs containing negative patterns. Using standard photolithography, silicon wafers were patterned with an SU-8 negative photoresist and subsequently silanized. A thick layer of polydimethylsiloxane (PDMS; Neyco, #RTV615) was then poured onto the wafer and cured at 70°C for at least 2 h. The cured PDMS layer was peeled off and used to generate epoxy replica molds.

PDMS devices were fabricated using standard soft lithography techniques as previously described [43]. Briefly, a 1:10 elastomer:curing agent mixture was poured into the custom-made epoxy molds, and cured at 70°C for 2 h. Following atmospheric plasma treatment for 15 s, PDMS chips bonded to 35-mm glass-bottom dishes (WPI, #FD35-100) and then incubated at 70°C for 15 min to reinforce the binding.

### Collagen gels

The collagen-cell mix was prepared on ice according to the provider’s instructions, using the following reagents: Collagen type-I rat tail solution (Corning, #354236), PBS (Euromedex, #EU1-2051-100), and 1 M NaOH (Fisher Scientific, #15673070). Briefly, a final volume, *V*_*f*_, of collagen-mix consisted of 0.36 × *V*_*f*_ of cells concentrated to 1 × 10^6^ cells/ml, the appropriate volume, *V*_*c*_, of collagen to reach the desired concentration, 0.1 × *V*_*f*_ of PBS 10x, 0.025 × *V*_*c*_ of 1 M NaOH, complemented with medium to reach the final volume *V*_*f*_. The final cell concentration in the mix was 0.3 − 0.35 × 10^6^ cells/ml, which avoided interaction due to self-generated gradients [44, 45].

After addition of NaOH, the mix was loaded in the PDMS chip (height of 350 *µ*m), and directly incubated at 37°C and 5% CO2 for 15 min. Collagen polymerization was stopped by submerging the chip with medium. Subsequently, cells were allowed to settle at 37°C and 5% CO2 for up to 2 h before starting imaging. After recording the random migration for at least 1 h, a chemokine was added to the surrounding medium to induce a chemotactic response. For DCs, CCL19 (Fisher Scientific, #17821413) was added to a final concentration of 40 nM, while CXCL2 (Biotechne, #452-M2-050/CF) was added to obtain a final concentration of 10 nM for neutrophil chemotaxis. The DC response was stable over several hours, while the neutrophil response was shorter due to receptor desensitization [46, 47], and we, thus, restricted the analysis to the time window where a response was observed.

### 2D devices

For surface functionalization, the PDMS device was plasma activated for 2 min and incubated with fibronectin (10 *µ*g/ml, Sigma-Aldrich, #F1141) at RT for 30 min. Subsequently, PDMS devices were washed with PBS three times, then incubated with medium (containing drugs where applicable) for 2 h at 37^°^C and 5% CO2 before cell loading. Cells labelled with Hoechst (Fisher Scientific, #12303553) were loaded at a concentration of 20 × 10^6^ cells/ml and allowed to settle and enter the chip for at least 2 h before addition of chemokine.

### Immunofluorescence

Samples were fixed with 4% paraformaldehyde (PFA; Fisher Scientific #11400580) for 30min at 37°C; then washed with PBS three times; To permeabilize cells, the sample was incubated with 0.1% Triton X-100 (Merck, #X100-100ML) for 10 min at room temperature (RT), subsequently washed with PBS three times, and blocked with PBS-2% BSA (Sigma-Aldrich, #A2153) overnight at 4^°^C. The fixed cells were incubated with Alexa Fluor 647 phalloidin (Thermo Fisher Scientific, #A22287), anti–*α*-tubulin antibody conjugated with fluorescein isothiocyanate (FITC) (abcam, #ab64503), and DAPI (Invitrogen, #D1306) or Hoechst 34580 (Thermo Fisher Scientific, #H21486) overnight at 4°C in a buffer containing PBS-2% BSA-0.05% saponin. Images were taken with a spinning-disc confocal microscope, with a 63X oil objective. The spinning disc-confocal microscope had a Yokogawa CSU-X1 spinning-disc head on a DMI-8 Leica inverted microscope equipped with a Hamamatsu OrcaFlash 4.0 Camera, a NanoScanZ piezo focusing stage (Prior Scientific) and a motorized scanning stage (Marzhauser). Z stacks of 40 slices with a spacing of 0.5 µm were taken. For images presented in Fig.5, z max projection is presented. The microtubule signal in collagen gels in Fig. 5f was enhanced by applying the DenoiSeg function of the CSBDeep plugin in Fiji.

### Live imaging

Migrating cells were imaged for 6 h with a 10x dry objective on a DMi8 inverted microscope (Leica), controlled by MetaMorph software (Molecular Devices). An on-stage incubation chamber maintained the temperature at 37^°^C and CO2 concentration at 5%. An image was taken every 30 s. Phase contrast was used for imaging cells migrating in collagen gels,

### Image processing and analysis

Image processing was performed using a custom made script in Fiji (Image J) software as detailed in [16]. The tiff stack was registered using the Image Stabilizer plugin, and the median image was subtracted. Next, a mean filter was applied to the entire stack and the result subtracted from the stack. Subsequently, a Gaussian blur filter was applied to the processed stack.

### Analysis of cell trajectories

Cells were then tracked using trackpy [48]. The trajectory analysis was restricted to the area closest to the chemokine source, spanning from the edge up to 600 µm. Furthermore, cells that covered a distance smaller than 20 µm, or had an average speed less than 0.5 µm were considered immobile, and removed from the analysis.

The directionality is calculated as cos *θ* = *v*_*x*_*/v*, the angular velocity is calculated *ω* = Δ*θ/*Δ*t*. To obtain the relative speed change, Δ*v/v*, the speed difference, Δ*v* is calculated as the difference between the speed when the cells are moving up the chemokine gradient, (*v*_cos *θ*=1_, and the speed when the cells are moving down the chemokine gradient, (*v*_cos *θ*=−1_. The relative speed change is then defined as (*v*_cos *θ*=1_ − *v*_cos *θ*=−1_)*/v*_cos *θ*=1_.

The conditional probability, *P* (*ω*|*θ*), shown in Fig. 3 was calculated from the joint probability distribution, *P* (*ω* ∩ *θ*), and the distribution of orientations, *P* (*θ*), according to *P* (*ω* ∩ *θ*)*/P* (*θ*) (see Supp Fig. 3 for *P* (*ω* ∩ *θ*)).

The predicted distribution of orientations *P* (*θ*) in Fig. 4 was obtained by integrating Eq.(2) using the Simpson’s rule in scipy with the experimentally obtained standard deviation of the turning angle as 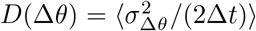. As the linear relationship between *ω* and *θ* is mostly restricted to the range [−*π/*2, *π/*2], the normalization constant was not analytically calculated but obtained by fitting the result to the experimentally obtained *P* (*θ*) using the curve fit function in scipy. Fitting does not change the shape of the predicted *P* (*θ*).

### Statistical analysis of experimental data

Statistical analysis of experiments was performed using Python 3 (NumPy and SciPy libraries) Line plots with shading represent the mean *±* SD of independent experiments. The number of independent experiments, N, or in the case of single representative experiments, the number of independent trajectories, n, is provided in the figure legends of the figure panel corresponding to the data. Statistical significance was determined by two-tailed unpaired or paired Student’s t test, where significance is reported as follows: ns: non-significant, *: p≤ 0.05, **:p ≤ 0.01, ***: p ≤ 0.001.

**Supp. Fig. 2.**
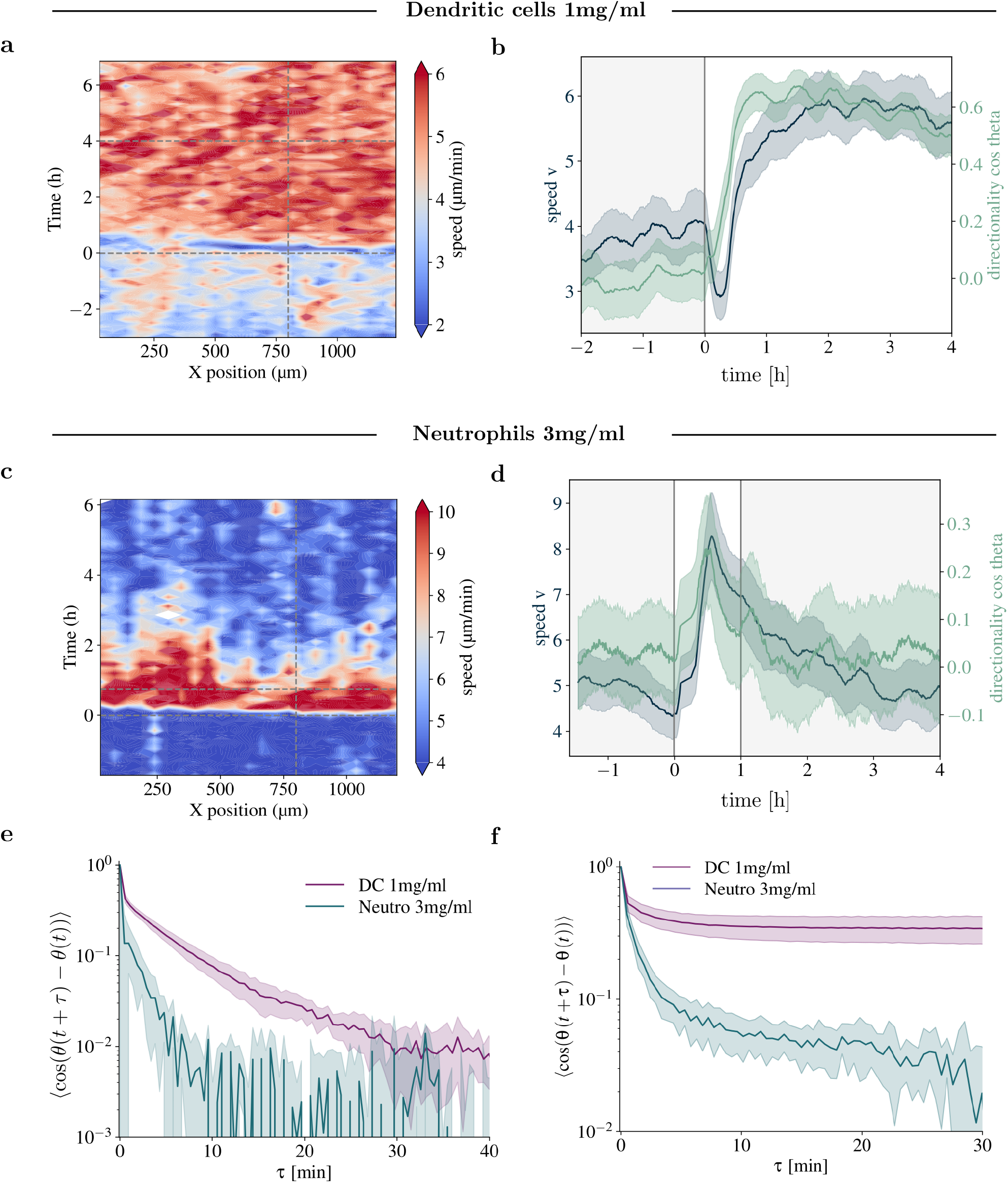
Chemotaxis of dendritic cells (DC) and neutrophils in porous collagen environements. **a** kymogrpah of the median speed and **b** averaged speed and directionality over time of dendritic cells in 1mg/ml collagen gel, **c** kymograph of the median speed and **d** averaged speed and directionality of neutrophils in 3mg/ml collagen gel, **e** directional autocorrelation in the absence of a chemokine gradient **f** directional autocorrelation in the presence of a chemokine gradient.

**Supp. Fig. 3.**
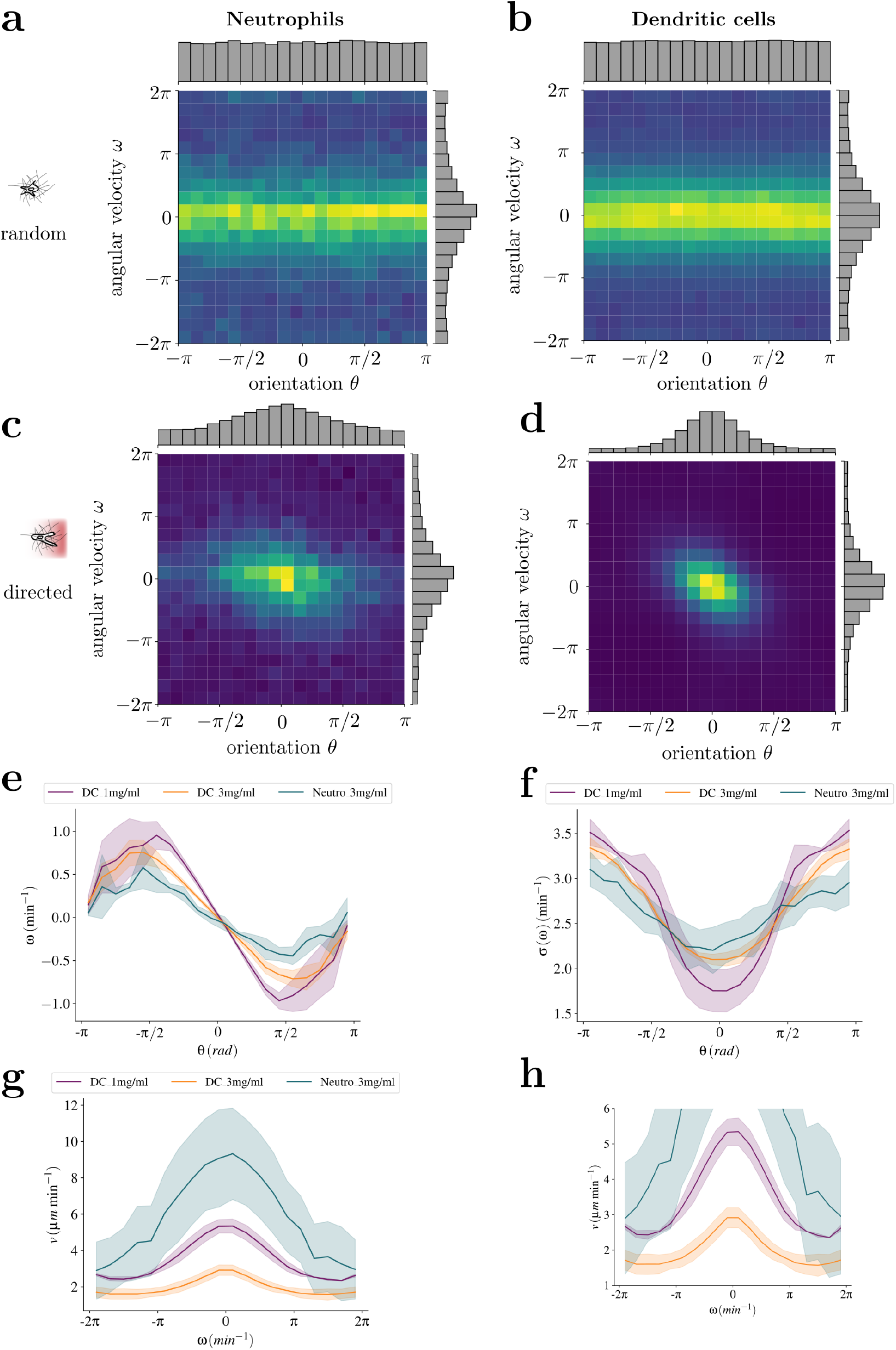
Torque and coloured noise. **a-b**, The joint probability distribution *P* (*ω* ∩ *θ*) with the individual distributions, *P* (*ω*) and *P* (*θ*), shown on the axis for neutrophils and DCs in the absence of a chemokine gradient; and **c-d** in the presence of a chemokine gradient. The conditional probability, *P* (*ω* | *θ*), in the main text is calculated according to *P* (*ω* ∩ *θ*)*/P* (*θ*). **e**, median angular velocity **f**, standard deviation of the angular velocity **g**, speed as a function of the angular velocity **h**, zoomed-in version of **g**. The line plot represent the mean ± SD of 5 independent experiments.

**Supp. Fig. 4.**
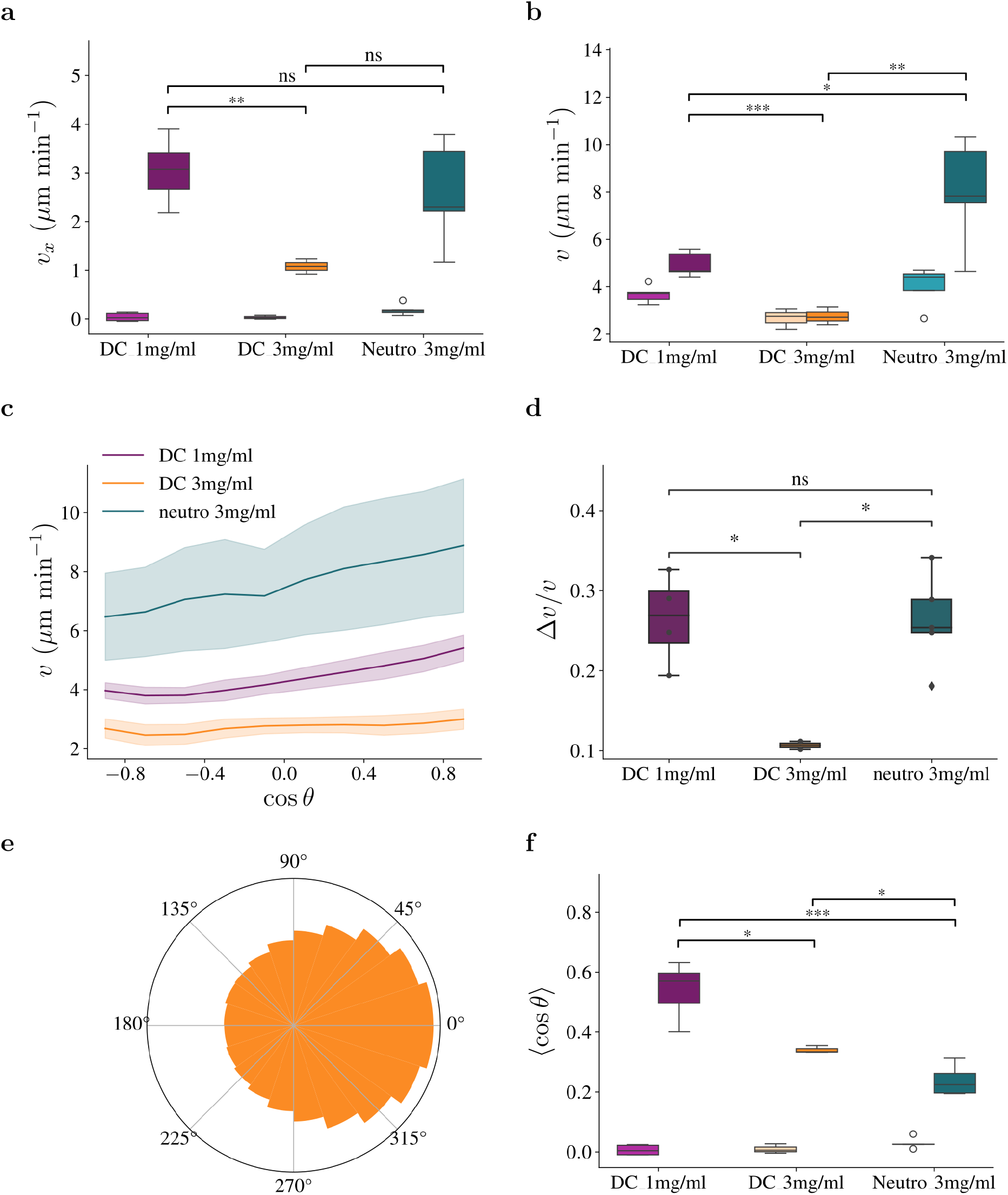
Effect of collagen concentration on chemotaxis of dendritic cells. **a**, Speed along the gradient direction, *v*_*x*_, **b**, Mean speed *v*, **c**, Speed as a function of the orientation towards the chemical gradient, cos *θ*, **d**, Relative speed change, (*v*_cos *θ*=1_ − *v*_cos *θ*=−1_)*/v*_cos *θ*=1_, **e**, Distribution of orientations *θ* for DCs migrating in 3mg/ml collagen gel, **f**, Directionality, cos *θ* = *v*_*x*_*/v*.

**Supp. Fig. 5.**
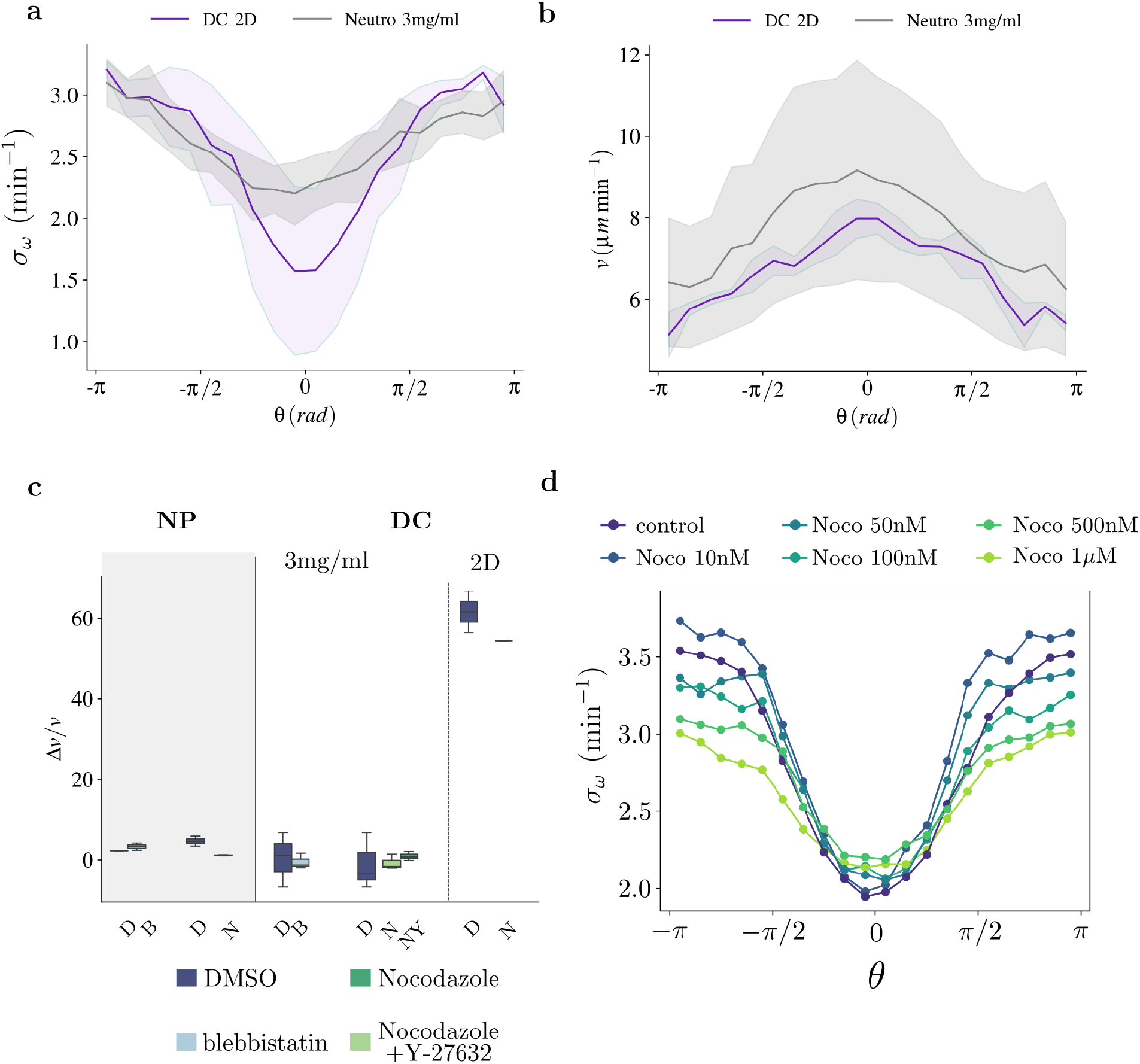
DC migration in 2D ressembles neutrophil migration in collagen gels. **a**, Standard deviation of the angular velocity as a function of the orientation **b**, Speed as a function of the angular velocity **c**, Effects of environment and drug treatments on mean speed **d**, nocodazole dose effect on the noise in angular velocity.

**Supp. Fig. 6.**
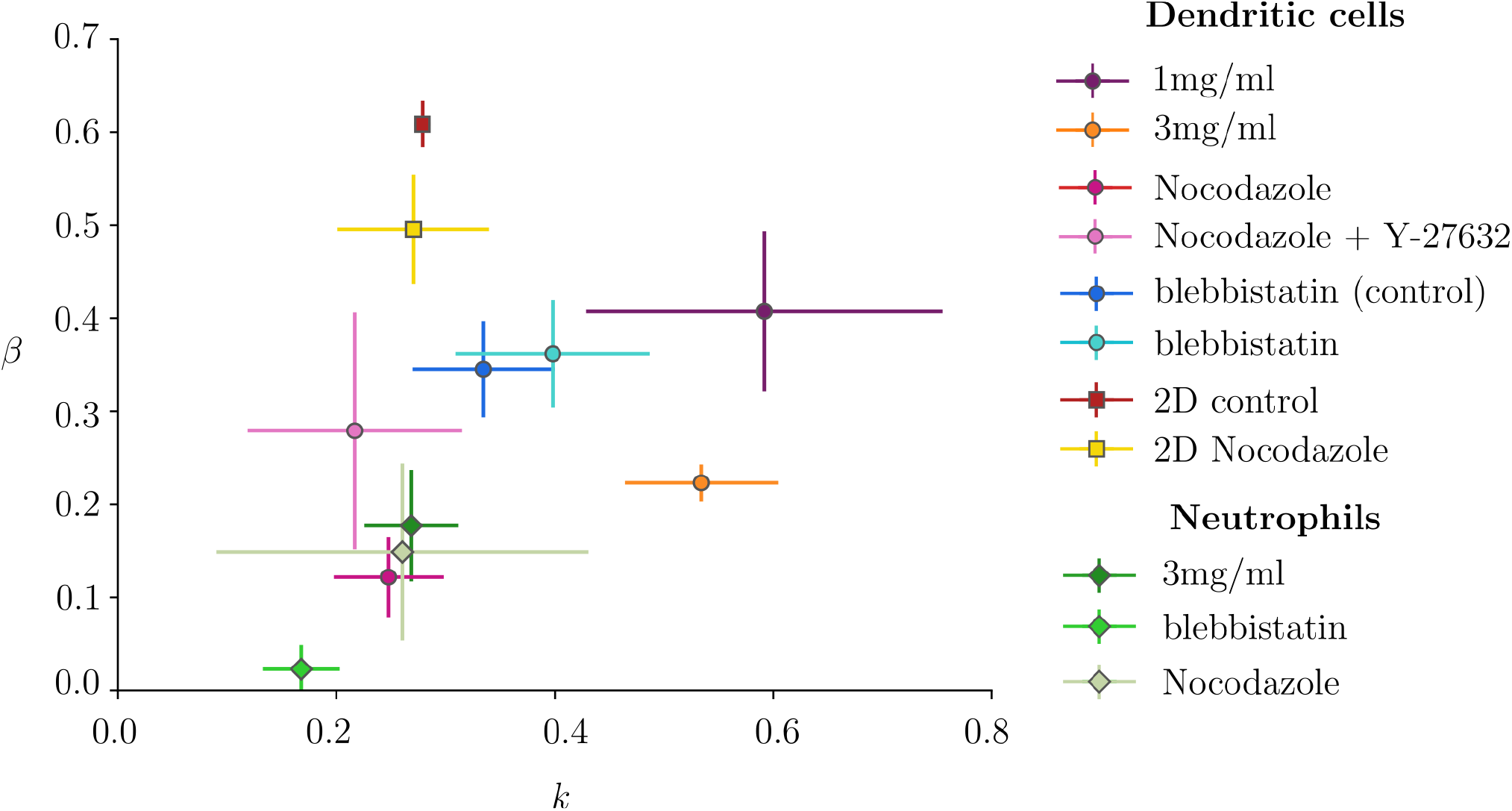
Chemotactic phase space. defined by the extent of noise in rotational diffusion, *β*, and the torque parameter *k*, without the scaling in the main text.

**Supp. Fig. 7.**
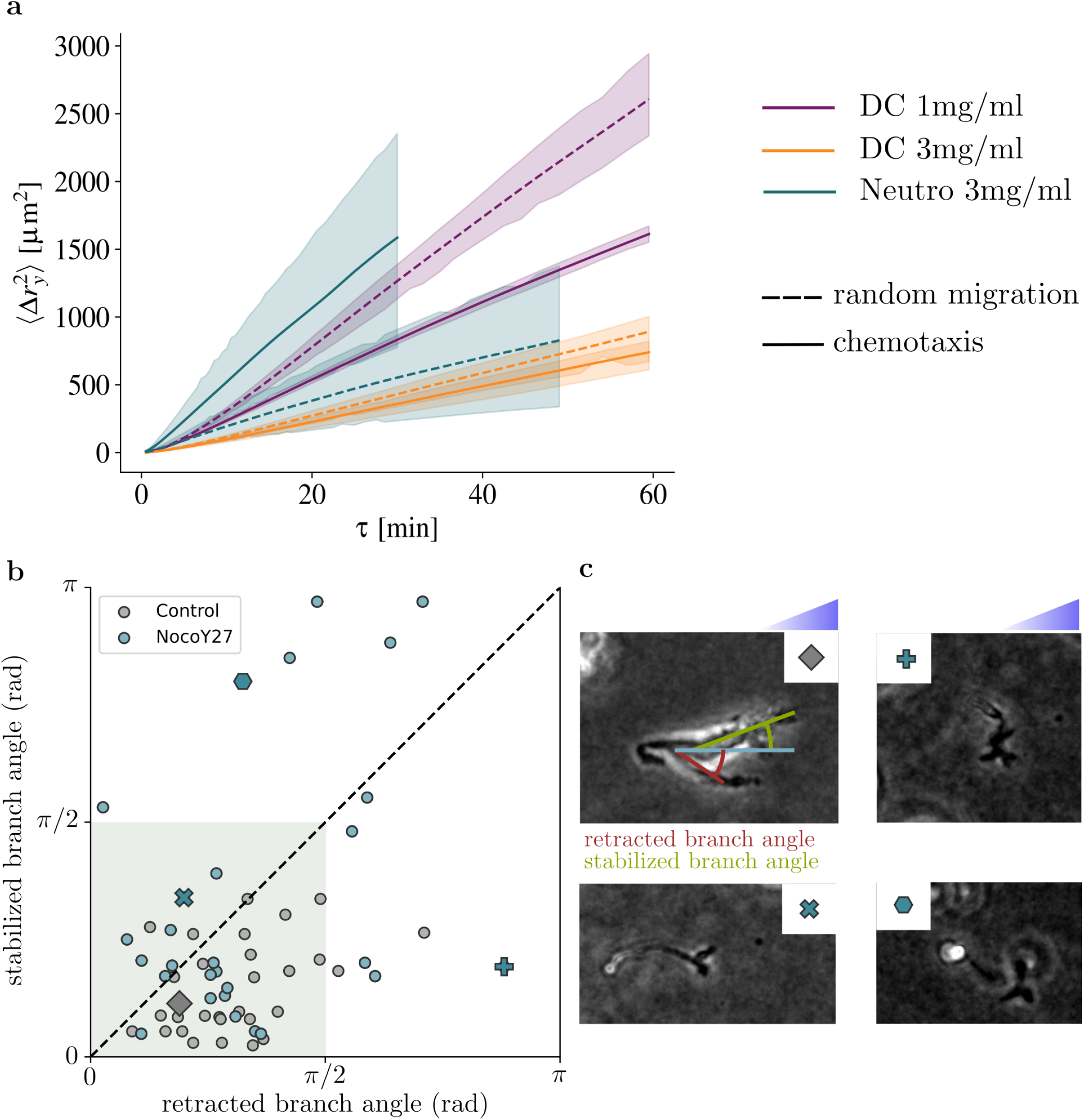
Microtubules determine the choice of branches in 3D migration. **a**, Mean squared displacement in the direction perpendicular to the chemokine gradient. Dashed lines: absence of a chemokine gradient, solid lines: presence of a chemokine gradient. **b**, Angles of retracted and stabilised branch during a branching decision in a 3mg/ml collagen gel in the presence of a CCL21 chemokine gradient for DCs treated with DMSO (grey data points) or nocodazole and Y27 (blue data points). The shaded box illustrates the range where both protrusions are formed up the gradient. If points are below the dashed 45^°^ line, the protrusion with the smaller angle is stabilised **c**, Examples of cells corresponding to data points shown in **b**.

