## Supplementary model information for "Torque-based immune cell chemotaxis in complex environments"

### Supplementary Material: Torque based immune cell chemotaxis in complex environments

Let us consider an Active Brownian Particle (ABP) as our model particle. In the standard case, an ABP performs a persistent random walk, moving at constant speed $v$ in direction $\theta$, which is subject to random white noise. At the single particle level, the dynamics of an ABP at position $\mathbf{x}$ can be described by the Langevin equation

$$\begin{aligned} \dot{\mathbf{x}} &= v\,\mathbf{e}(\theta) \\ \dot{\theta} &= \sqrt{2D_r}\xi(t), \end{aligned} \tag{1}$$

where dot denotes the time derivative, unit vector $\mathbf{e} = [\cos\theta, \sin\theta]$. The white noise in Eq. (1) obeys $\langle \xi_r(t) \rangle = 0$ and $\langle \xi_r(t)\xi_r(t') \rangle = \delta(t - t')$.

To examine chemotactic strategies, we consider the case of an ABP responding to a non-homogenous field of attractant, whose concentration is increasing along the $x$-direction. Thus, a chemotactic strategy needs to introduce a drift in $x$. Two parameters in Eq. (1) can be modified to achieve this drift in response to a changing attractant concentration: the speed, $v$, and the persistence time, $1/D_r$.

In order to perform chemotaxis, particles have to measure attractant concentrations. The physical limits of this based on particle size have been discussed previously, e.g., in [1]. In the following, we will assume that the particle is able to perform an estimate of either a) the attractant concentration, b) a projection of the attractant gradient along a major axis, or c) the complete gradient. The ability of a particle to perform these estimates is related to its size and shape [2], as illustrated in figure Fig. 1. In the following, we will discuss the different strategies of biasing the motility parameters which follow from the gradient estimated used.

#### 1 Point particle

The simplest chemotaxis strategy involves small, point-like particles. These particles are too small to perform a measurement of the gradient, but can sense the chemical concentration locally. In a non-uniform concentration field, particles then adjust their motility parameters based on these local estimates. Eq. (1) is modified to have the parameters speed and/or orientation as a function of the particle position $\mathbf{x}$

$$\begin{aligned} \dot{\mathbf{x}} &= v(\mathbf{x})\mathbf{e}(\theta) \\ \dot{\theta} &= \sqrt{2D_r(\mathbf{x})}\xi_r(t), \end{aligned} \tag{2}$$

with the corresponding Fokker-Planck equation

$$\frac{\partial P(\mathbf{x},\theta,t)}{\partial t} = -\nabla\left[v(\mathbf{x})\mathbf{e}P\right] + \frac{\partial^2}{\partial\theta^2}\left[D_r P\right]. \tag{3}$$

Following the derivation presented in [3, 4], the microscopic system can be approximated by a drift-diffusion equation

$$\partial_t\varphi = -\nabla(-D\nabla\varphi + \mathbf{V}\varphi), \tag{4}$$

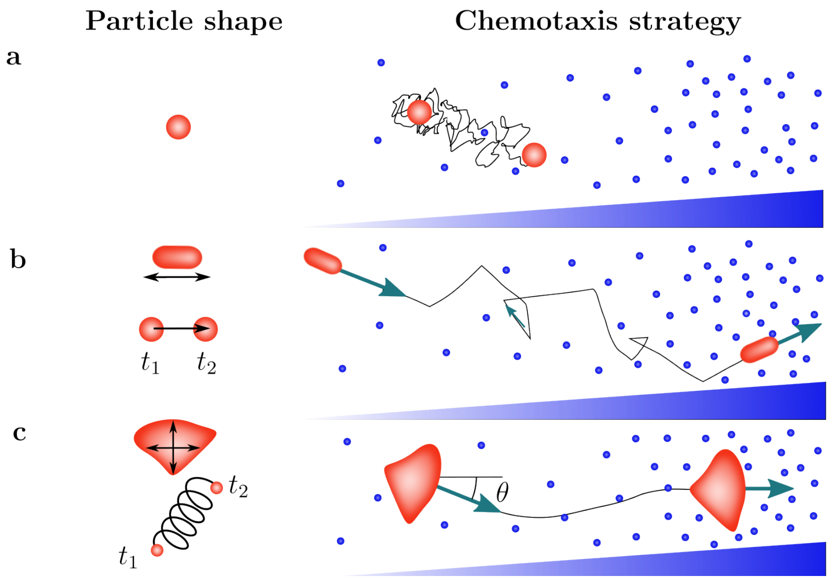


**Fig. 1 Gradient sensing and chemotactic strategy depend on particle shape.** **a**, Point-like particle employing a local response; **b**, Chemotactic response based on $\cos\theta$, which is determined either by direct sensing along the length of a polar particle or approximated by memory of measurements at different time points; **c**, Biasing the orientation towards a gradient $\theta$, which is measured either directly along the axes of an extended particle or approximated, e.g., along fixed helical trajectories.

where

$$D = \frac{v^2}{d(d-1)D_r}, \tag{5}$$

and

$$\mathbf{V} = \frac{-v\nabla v}{d(d-1)D_r}. \tag{6}$$

In a bounded domain, we can obtain a steady-state solution with $\nabla J = 0$. Introducing an effective potential

$$\psi(\mathbf{x}) = \int dq \frac{\mathbf{V}(q)}{D(q)} = -\int dq \frac{\nabla v(q)}{v(q)}, \tag{7}$$

we find that only a spatially-varying speed, $v(\mathbf{x})$, has an effect on the steady-state distribution

$$\varphi_s(\mathbf{x}) \propto \exp(-\psi(\mathbf{x})), \tag{8}$$

which has also been shown for run-and-tumble particles [5]. Thus, to achieve chemotaxis towards a source, the particle has to slow down upon approaching the source. This possible approach to bias the steady state distribution of cells based on the local speed has previously been discussed as a chemotaxis strategy [5, 6]. The effect of a spatially varying reorientation term, on the other hand, is generally considered negligible. These conclusions are based on the shape of Eq. (8), which is a steady-state ensemble average over the population. Alternatively, we could consider the local drift experienced by a single particle to analyse the dynamics. For the initial condition $x(0) = 0$, $\psi(x,t)$ is the single cell propagator, which defines the drift

$$\partial_t \langle x \rangle = \int_{-\infty}^{\infty} x \partial_t \varphi(x,t) dx. \tag{9}$$

Inserting Eq. (4) in Eq. (9) and applying integration by parts twice yields

$$\begin{aligned}\partial_t \langle x \rangle &= \int_{-\infty}^{\infty} x\nabla(D\nabla\varphi)dx - \int_{-\infty}^{\infty} x\nabla(\mathbf{V}\varphi)dx \\ &= \langle \nabla D \rangle + \langle V \rangle,\end{aligned} \tag{10}$$

where we used $\varphi(-\infty, t) = \varphi(\infty, t) = 0$. If the speed is constant, i.e. $v(x) = v_0$, this simplifies to

$$\partial_t \langle x \rangle = \langle \nabla D \rangle = -\frac{v_0^2}{d(d-1)} \langle \frac{D_r'(x)}{D_r(x)^2} \rangle, \tag{11}$$

and shows that a position-dependent noise term, $D_r(x)$, indeed generates a local drift. Taking $D_r(x) = \text{const}$ and $v(x)$ in Eq. (10), $\partial_t \langle x \rangle \propto -v\nabla v$, leads to the counter-intuitive observation that the particle experiences a drift towards regions with a larger speed, which is the opposite result of the steady-state observable. In conclusion, a basic chemotaxis strategy is to modulate speed and directional persistence in different attractant concentrations, but the choice of observable is crucial to evaluate the strategies.

### 2 Polar particle

In the previous section, the parameters $v$ and $D_r$ were functions of $\mathbf{x}$, i.e. the parameters are modified based on local estimates of the chemical concentration. In this section, we turn to particles which can estimate the gradient along their moving direction, either directly by measuring along the polar axis of the cell body or indirectly by memory. As a result, the parameters are a function of the orientation of the particles, $\theta$. Note that measuring the change in chemical concentration along the particle body or travelled direction means that the particle is effectively measuring a projection of the gradient, $\cos\theta$. As a result, a particle cannot distinguish between an increase in gradient strength and an improved alignment with a (constant) gradient.

#### 2.1 Variable rotational diffusion

For simplicity, we first consider the case of a variable rotational diffusion coefficient and constant speed, i.e. $D_r(\theta) = f(\theta)$ and $v(\theta) = \text{const}$. Without ambiguity in the stochastic notation, the experimental data on the distribution of orientations, shown in the main text, can be modelled by the Fokker-Planck equation

$$\partial_t P(\theta, t) = -\partial_\theta[\mu(\theta)P(\theta, t)] + \partial_\theta^2[D(\theta)P(\theta, t)], \tag{12}$$

where $\mu(\theta)$ and $D(\theta)$ are the drift and (biased) diffusion term, respectively. In the case of a chemotactic strategy based on the modulation of angular noise, we have $\mu(\theta) = 0$ and $D(\theta) = D_r(\theta)$. To make the link with a microscopic model as in Eq. (1), we need to choose a Langevin equation. The presence of multiplicative noise in stochastic processes requires to choose an interpretation of the stochastic noise to define the stochastic integral, which is used to derive the corresponding FPE from the Langevin equation. While other conventions could be possible, we choose a Langevin equation with Ito convention as this convention is consistent with the FPE in Eq. (12), imposed unambiguously by the data. The corresponding Langevin equation is, thus,

$$\begin{aligned}\theta &= v(\theta)\mathbf{e}(\theta) \\ \dot{\theta} &= \sqrt{2D_r(\theta)}\xi_r(t),\end{aligned} \tag{13}$$

where $\langle \xi_r(t) \rangle = 0$ and $\langle \xi_r(t)\xi_r(t') \rangle = \delta(t - t')$. The FPE for the stochastic process given by Eq. (13) is

$$\partial_t P(\mathbf{x}, \theta, t) = -v\nabla[\mathbf{e}P] + \partial_\theta^2[D_r(\theta)P], \tag{14}$$

where the multiplicative noise introduces an effective drift required for chemotaxis. To obtain the FPE given in the main text, we simplify (2). The dynamics of $\theta$ do not depend on $x$ and, for large enough $\omega$, it should relax quickly to a quasi-stationary state in which $d\theta/dt \to 0$, and $P(\theta, t) = P_s(\theta)$. We thus assume a separation of time-scale so that

$$P(\mathbf{x}, \theta, t) = g(\mathbf{x}, t_1) f(\theta, t_0), \tag{15}$$

where $t_0 = t$ and $t_1 = \epsilon t$ are the fast and slow time scale, respectively, and $d/dt = \partial/\partial t_0 + \epsilon \partial/\partial t_1$. Thus, inserting into Eq. (14) we get

$$(\epsilon \partial_{t_1} g(\mathbf{x}, t_1)) f + g \partial_{t_0} f(\theta, t_0) = -\epsilon v \nabla \left[ \mathbf{e} g(\mathbf{x}, t_1) f(\theta, t_0) \right] + g(\mathbf{x}, t_1) \partial_\theta^2 [D_r(\theta) f(\theta, t_0)]. \tag{16}$$

For $\epsilon \ll 1$, this simplifies to

$$\partial_{t_0} f(\theta, t_0) = \partial_\theta^2 [D_r(\theta) f(\theta, t_0)] = -\partial_\theta J_{t_0}, \tag{17}$$

where we identified the flux $J_{t_0} = -\partial_\theta [D_r(\theta) f(\theta, t_0)]$ and recovered the FPE given by Eq. (12). This is a diffusion process in 1D with periodic boundary conditions $[a, b] = [-\pi, \pi]$ and state-dependent diffusion term. As $D_r(\theta)$ is a periodic function, we can set $J = 0$ [7]. Thus, the stationary distribution in angle $\theta$ is given by

$$P_s(\theta) = \frac{\mathcal{N}}{D_r(\theta)}, \tag{18}$$

where $\mathcal{N}$ is a normalization factor to ensure $\int P_s(\theta) \mathrm{d}\theta = 1$. At long times, i.e. $\epsilon \gg 1$, Eq. 16 simplifies to

$$\partial_{t_1} g(\mathbf{x}, t_1) f_s(\theta) = -v \nabla \left[ \mathbf{e} g(\mathbf{x}, t_1) f_s(\theta) \right]. \tag{19}$$

Multiplying by $\mathbf{x}$, averaging over $\theta$ and integrating over time, we obtain

$$\langle y \rangle = t\, v_0 \int_{-\pi}^{\pi} \mathrm{d}\theta P_s(\theta) \sin\theta = t\, v_0 \, \langle \sin\theta \rangle, \tag{20}$$

and

$$\langle x \rangle = t\, v_0 \int_{-\pi}^{\pi} \mathrm{d}\theta P_s(\theta) \cos\theta = t\, v_0 \, \langle \cos\theta \rangle, \tag{21}$$

which is the chemotactic drift as a function of the directionality, $\langle \cos\theta \rangle$.

In the main text, we consider the bias in the angular noise given by

$$D_r(\theta) = D_r^0 (1 - \beta \cos\theta)^2, \tag{22}$$

which gives

$$P_s(\theta) = \frac{\mathcal{N}}{D_r(\theta)} = \frac{\sqrt{(1-\beta^2)^3}}{2\pi(-1+\beta\cos\theta)^2}. \tag{23}$$

Note that due to normalization, $P_s(\theta)$ does not depend on the base-line diffusivity $D_r^0$. The directionality is thus

$$\langle \cos\theta \rangle_p(\beta) = \int_{-\pi}^{\pi} P_s(\theta) \cos(\theta) \mathrm{d}\theta = \beta. \tag{24}$$

and the chemotactic drift along the chemotactic gradient follows as $\langle x \rangle = tv \, \langle \cos\theta \rangle$.

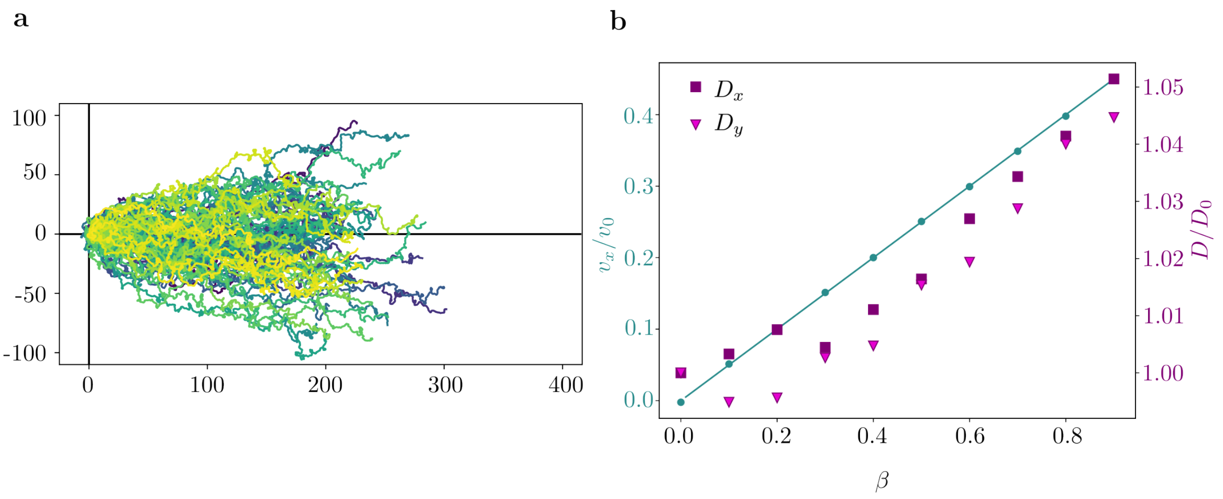


**Fig. 2 Chemotactic response of a particle with speed modulation. a** Simulated trajectories for a polar particle with (non-dimensionalized) $v(\theta) = (1 - \beta \cos\theta)$, where $\beta = 0.7$, leading to a chemotactic drift of $v_x/v_0 = 0.35$. **b** Drift velocity $v_x/v_0$ and diffusion coefficients $D_x$, $D_y$ as a function of the biasing parameter $\beta$ (symbols, simulations; lines, analytical results).

#### 2.2 Variable speed

A successful chemotactic response can also be based by modulating the speed as a function of the projected gradient, $\cos\theta$. For simplicity, assume $D_r(\theta) = \text{const}$ and varying speed $v(\theta)$. The average position over time then follows

$$\langle x \rangle = t \int_{-\pi}^{\pi} \mathrm{d}\theta \, v(\theta) \cos(\theta) P_s(\theta) \tag{25}$$

In this case, $P_s(\theta)$ is that of a Gaussian process, i.e. in periodic boundary conditions the distribution is uniform, and normalised by the length of domain, i.e. $P_s(\theta) = 1/2\pi$. Indeed we show in the main text that neutrophils and DCs in 1mg/ml modulate their speed in response to a gradient, in line with previous reports, e.g., on neutrophils in zebrafish [6]. The data suggests that the speed depends linearly on $\cos\theta$, e.g.,

$$v(\theta) = v_0(1 + \beta \cos\theta), \tag{26}$$

where $v_0$ is the speed at $\theta = \pm\pi/2$. For this speed modulation we obtain the chemotactic drift as

$$\langle x \rangle = \frac{v_0 \, \beta}{2} \, t, \tag{27}$$

and, as expected, $\langle y \rangle = 0$. In this case, the maximal achievable relative drift speed is thus $v_x/v_0 = 0.5$ with $\beta = 1$. As shown in Fig. 2, speed modulation leads to clear a chemotactic bias. The diffusion coefficients in direction perpendicular and parallel to the chemotactic gradient increase with only marginally increasing alignment parameter $\beta$ for the linear speed modulation given by Eq. (26). Other speed modulations are possible and can lead to a stronger increase in both diffusion coefficients but are not suggested by the data for either neutrophils or DCs.

### 3 Extended particle

Finally, if the particle is able to estimate the gradient not just along the polar axis but also the lateral axis, it may directly turn towards the direction of the gradient, with an aligning torque acting on the orientation vector. The angular dynamics can then be described by an Ornstein-Uhlenbeck process under periodic boundary conditions as follows

$$
\begin{aligned}
\theta &= v_0 \mathbf{e}(\theta) \\
\dot{\theta} &= -k\theta + \sqrt{2D_r}\xi_r(t),
\end{aligned}
\tag{28}
$$

where $\kappa$ is an effective torque term. We can identify the characteristic time and length scales as

$$
t_c = \frac{1}{2D_r} \quad \text{and} \quad x_c = \frac{v_0}{2D_r}. \tag{29}
$$

Using these transformations, the dimensionless Langevin system is

$$
\begin{aligned}
\dot{\mathbf{x}} &= \mathbf{e}(\theta) \\
\dot{\theta} &= -\kappa\theta + \xi_r(t),
\end{aligned}
\tag{30}
$$

where the Gaussian process was transformed as $\xi_r(\tilde{t}) = \sqrt{1/t_c}\,\xi_r(t)$ and $\kappa = k/2D_r$. The corresponding Fokker-Planck equation is

$$
\partial_t P(\mathbf{x}, \theta, t) = -\nabla\left[\mathbf{e}P\right] + \frac{1}{2}\partial_\theta^2 P - \partial_\theta\left[\kappa\theta P\right]. \tag{31}
$$

Again, the angle does not depend on $x$ and, for large enough $\kappa$, it should relax quickly to a quasi-stationary state in which $d\theta/dt \to 0$, and $P(\theta, t) = P_s(\theta)$. The corresponding Fokker-Planck equation follow as

$$
\partial_t P(\theta, t, |\theta_0, t_0) = \frac{1}{2}\partial_\theta^2 P + \partial_\theta(\kappa\theta P) = \partial_\theta J, \tag{32}
$$

where we introduced the probability flux $J$. In stationary state $\partial_t P = 0$ and, thus, $\partial_\theta J = 0$. According to Risken, if we do not distinguish whether a full rotation has been made or not, the stationary probability distribution should be periodic, and $J$ is determined by the periodicity condition. These periodic boundary conditions can only be fulfilled if the drift and diffusion coefficients are also periodic with period $L = 2\pi$. The stationary probability distribution under periodic boundaries $[a, b]$ is given as

$$
P_s(\theta) = \frac{\mathcal{N}}{B(\theta)} \exp\left(2\int_a^\theta \frac{A(\theta')}{B(\theta')} d\theta'\right), \tag{33}
$$

where $A(\theta)$ and $B(\theta)$ are the drift and diffusion terms in the Fokker-Planck equation, respectively. For $A(\theta) = \omega\theta$ and $B(\theta) = -1$, this results in

$$
P_s(\theta) = \mathcal{N}\exp(-\kappa(\theta^2 - a^2)), \tag{34}
$$

where the normalization constant $\mathcal{N}$ is determined as

$$
\mathcal{N} = \sqrt{\pi/\kappa}\exp(\pi^2\kappa)\mathrm{erf}(\pi\sqrt{\kappa}). \tag{35}
$$

Then, the directionality is

$$
\langle\cos\theta\rangle = \int_{-\pi}^{\pi} \mathrm{d}\theta P_s(\theta)\cos\theta = \frac{\exp(-1/4\kappa)}{\mathrm{erf}(\pi\sqrt{\kappa})}\mathrm{Im}\left\{\mathrm{erfi}\left(\frac{1 + 2i\pi\kappa}{2\sqrt{\kappa}}\right)\right\}, \tag{36}
$$

and the dimensionless chemotactic drift follows as $\langle x\rangle = t\,\langle\cos\theta\rangle$.

##### 3.1 Equivalence of polar and extended particle processes

Both, a torque based strategy and a strategy based on a modulation of the rotational diffusion, introduce a drift term in the Fokker-Planck equation. This could raise the question if the two processes are equivalent, e.g., if there is a choice of $D_r(\theta)$ possible, which allows to derive an equivalent OU process.

$$\partial_t P(\theta, t, |\theta_0, t_0) = D_r \partial_\theta^2 P + \kappa\theta\partial_\theta P + \kappa P, \tag{37}$$

where $D_r$ is a constant. Comparing Eq. (37) and the Fokker-Planck of biased diffusion in Ito form

$$\partial_t P(\theta, t, |\theta_0, t_0) = D_r(\theta)\partial_\theta^2 P + 2\partial_\theta D_r(\theta)\partial_\theta P + \partial_\theta^2 D_r(\theta) P, \tag{38}$$

suggests that $D_r(\theta) = \frac{1}{2}\kappa\theta^2 + c$, where $c$ is a constant. However, this form of $D_r(\theta)$ introduces a term $\frac{1}{2}\kappa\theta^2\partial_\theta^2 P$ in the first term on the left hand side of Eq. (38), which cannot be matched with the OU description. In fact, any choice of convention (Ito, Stratonovich or anti-Ito) will introduce this term, and the two processes are thus not equivalent.

##### 3.2 Comparison polar and extended particle

To estimate the influence of the different strategies on the chemotactic response in the main text, the following ratio was solved numerically

$$\Psi(k', \beta) = \frac{\langle\cos\theta\rangle_t(k')}{\langle\cos\theta\rangle_t(k') + \langle\cos\theta\rangle_p(\beta)}. \tag{39}$$
